# Multivariate resilience indicators to anticipate vector-borne disease outbreaks: a West Nile virus case-study

**DOI:** 10.1101/2024.12.09.627472

**Authors:** Clara Delecroix, Quirine ten Bosch, Egbert H. Van Nes, Ingrid A. van de Leemput

**Affiliations:** Environmental Sciences Group, Wageningen University & Research, the Netherlands; Infectious Disease Epidemiology, Wageningen University & Research, the Netherlands

## Abstract

**Background and aim:** To prevent the spread of infectious diseases, successful interventions require early detection. The timing of implementation of preventive measures is crucial, but as outbreaks are hard to anticipate, control efforts often start too late. This applies to mosquito-borne diseases, for which the multifaceted nature of transmission complicates surveillance. Resilience indicators have been studied as a generic, model-free early warning method. However, the large data requirements limit their use in practice. In the present study, we compare the performance of multivariate indicators of resilience, combining the information contained in multiple data sources, to the performance of univariate ones focusing on one single time series. Additionally, by comparing various monitoring scenarios, we aim to find which data sources are the most informative as early warnings.

**Methods and results:** West Nile virus was used as a case study due to its complex transmission cycle with different hosts and vectors interacting. A synthetic dataset was generated using a compartmental model under different monitoring scenarios, including data-poor scenarios. Multivariate indicators of resilience relied on different data reduction techniques such as principal component analysis (PCA) and Max Autocorrelation Factor analysis (MAF). Multivariate indicators outperformed univariate ones, especially in data-poor scenarios such as reduced resolution or observation probabilities. This finding held across the different monitoring scenarios investigated. In the explored system, species that were more involved in the transmission cycle or preferred by the mosquitoes were not more informative for early warnings.

**Implications:** Overall, these results indicate that combining multiple data sources into multivariate indicators can help overcome the challenges of data requirements for resilience indicators. The final decision should be based on whether the additional effort is worth the gain in prediction performance. Future studies should confirm these findings in real-world data and estimate the sensitivity, specificity, and lead time of multivariate resilience indicators.

**Author summary:** Vector-borne diseases (VBD) represent a significant proportion of infectious diseases and are expanding their range every year because of among other things climate change and increasing urbanization. Successful interventions against the spread of VBD requires anticipation. Resilience indicators are a generic, model-free approach to anticipate critical transitions including disease outbreaks, however the large data requirements limit their use in practice. Since the transmission of VBD involves several species interacting with one another, which can be monitored as different data sources. The information contained by these different data sources can be combined to calculate multivariate indicators of resilience, allowing a reduction of the data requirements compared to univariate indicators relying solely on one data source. We found that such multivariate indicators outperformed univariate indicators in data-poor contexts. Multivariate indicators could be used to anticipate not only VBD outbreaks but also other transitions in complex systems such as ecosystems’ collapse or episodes of chronic diseases. Adapting the surveillance programs to collect the relevant data for multivariate indicators of resilience entails new challenges related to costs, logistic ramifications and coordination of different institutions involved in surveillance.

## Introduction

To prevent the spread of infectious diseases, successful interventions require early detection. The timing of implementation of preventive measures is crucial, but as outbreaks are hard to anticipate, control efforts often start too late. The use of resilience indicators to anticipate infectious disease outbreaks has attracted attention in the past decade: they are generic, model-free indicators used to anticipate critical transitions in complex systems (1,2). These indicators typically rely on the theory of critical slowing down, stating that a system approaching a critical transition, such as the start of an epidemic, loses its resilience and recovers slower from external perturbations (3,4). In the context of infectious diseases, this translates into a longer time for minor outbreaks to resolve. This slow behaviour can be observed in trends of so-called resilience indicators, namely autocorrelation and variance. Such trends are usually calculated in the incidence time series using a rolling window. Resilience indicators have proven to be able to anticipate upcoming epidemics up to several months in advance for infectious diseases, including mosquito-borne diseases (1,5–7). However, the large data requirements limit their use in practice.

Surveillance of multi-host diseases is challenging to achieve, as many, sometimes unknown, species can participate in the transmission, which leaves choices on which sources to sample from (8). This is the case for West Nile virus for instance: the transmission burden can be monitored in mosquito pools by estimating the proportion of infected mosquitoes or the proportion of infected pools (9), but also by monitoring infections in humans, livestock, or wildlife through public health and veterinary surveillance systems (10). Choices then must be made on where to direct sampling efforts. Collecting data from multiple sources would either increase the sampling costs and efforts, or reduce the quality or sampling frequency of each data source. Contrarily, focusing on one single data source enables the relevant authorities to concentrate all their efforts on sampling one source meticulously, but the information contained in other sources may be missed. Additionally, when several species act as enzootic hosts and react differently to the disease, one can wonder which species is the most informative for early warnings of future outbreaks.

When data from multiple sources are available, the information they contain can be combined to calculate multivariate indicators of resilience (11). Previous research showed that multivariate indicators signal upcoming critical transitions in the same way univariate resilience indicators do, using simulated plant-pollinator data with a generic model (11). These multivariate indicators rely on data reduction techniques to combine multiple data streams, such as Principal Component Analysis (PCA) and Maximum Autocorrelation Factors (MAF). Common resilience indicators such as autocorrelation and variance are then computed in the obtained combined time series. Similar multivariate indicators have been investigated in simulated incidence time series of metapopulation models and could signal upcoming epidemics (12). However, they have not been compared to univariate indicators. As multivariate indicators require a greater monitoring effort, researching the best monitoring strategy would be of use to inform monitoring practices.

That question is not limited to infectious disease outbreaks: anticipating critical transitions in multivariate systems is relevant for a diversity of research areas, such as ecology (13,14) or finance (15). In many complex systems, several entities interact, leaving choices on where to direct monitoring efforts. These entities can interact heterogeneously via complex networks, and the interactions between such entities are not always well understood (16). Additionally, some entities seemingly affecting the dynamics of the system more directly are not always the easiest to monitor. For instance, food webs or ecosystems that are nearing collapse often contain many interacting species, for which central endangered species are not always easy to monitor (17). Previous research showed that some species can display more reliable (18) and earlier (19) signals using resilience indicators as early warning, but this has not been studied in epidemiology. Additionally, the prediction performance of multivariate and univariate indicators has never been compared.

West Nile virus (WNV) is a relevant example of a multi-host, vector-borne disease (VBD). WNV is transmitted between birds and mosquitoes, constituting the enzootic cycle. Humans and horses can get infected but do not transmit the virus onwards and are thus considered dead-end hosts. While human and horse cases can seem easy to monitor via public health and veterinary authorities, the small proportion of symptomatic cases results in low reporting probabilities ((10,20). On the other hand, monitoring wild birds is often done by collecting and analyzing dead birds (21). Monitoring live birds is also possible but requires a greater effort as they must be caught and sampled by bird ringers or citizen scientists, e.g. (22–24), and is thus more costly than sampling dead birds. Furthermore, the species contributing to the transmission of VBDs like WNV are not all known. Sometimes an unknown, hidden (i.e. not dying from the disease) reservoir can be central to the transmission dynamics (25). Infected mosquito pools can be monitored via traps, but estimating the prevalence of WNV in mosquitoes is complicated by logistic and cost limitations (26). Finally, as the transmission is dependent on the ecosystem setting, such as the abundance of the different hosts and vectors as well as vector feeding preferences, changes in ecosystems can impact the transmission if, e.g., several host species or vector species compete (27), making the task of identifying which species contribute most to transmission complex and context-specific. It is then relevant to know how to identify informative data sources to direct sampling efforts to anticipate future WNV outbreaks.

In the present study, we compare multivariate and univariate indicators of resilience under different monitoring scenarios as an early warning for upcoming epidemics of vector-borne diseases. We use WNV as a case study, as it affects multiple host and vector species, and can be monitored in numerous ways. A synthetic dataset is generated using a simple, well-studied WNV model. We explore several monitoring scenarios focusing on different data types, including data-poor scenarios with low reporting probability or resolution, and try to identify what makes a species more or less informative for early warnings.

## Materials and methods

### Model

We used a well-known compartmental Ross Macdonald model of WNV for which the dynamics are well-studied. The model is adapted from (28), and is used to generate time series. Our model considers the vector population and four host populations: two generic bird species acting as amplifying hosts, and humans and horses acting as dead-end hosts (Fig 1A). The two bird species differ in how they are affected by the disease. Bird species V is a visible, i.e. detectable, reservoir for the disease: 30% of the infected individuals die from infection (as can be the case for corvids, for instance (29)). Bird species H is a hidden reservoir: it is preferred by mosquitoes but it does not die from infection, hence complicating the detection of transmission in this species through dead bird surveillance.

**Fig 1.**
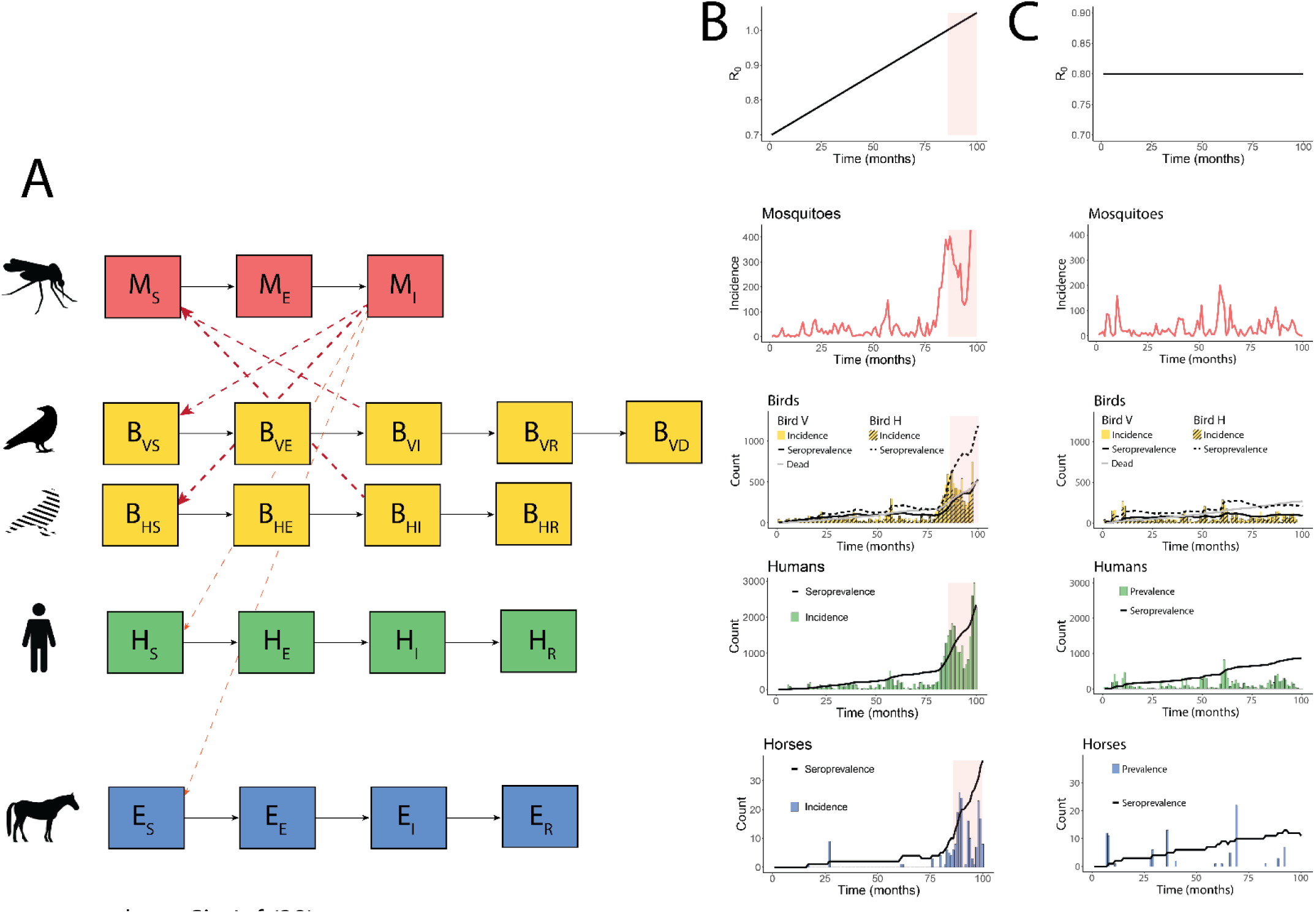
WNV model and simulated data (A) WNV compartmental model used in the analyses. (B) Example of simulated data with an upcoming epidemic, simulated by increasing R_0_ from 0.7 to 1. The shaded region indicates where R_0_ is above 1. (C) Example of simulated data with no upcoming epidemic, simulated with a stable R_0_=0.8. In (B) and (C), the time series were aggregated per month to improve the readability of the figure.

The mosquito population was subdivided into susceptible mosquitoes *M_S_*, exposed mosquitoes *M_E_*, and infectious mosquitoes *M_I_*. Mosquitoes get infected with a probability *p_M_* when biting infectious birds, which they encounter with a probability *δ_M_k*, with *δ_M_* denoting the fraction of non-diapausing mosquitoes and *k* the biting rate of mosquitoes. Additionally, we introduced biting preference coefficients *p_i_* for each host species: bird H was the preferred host, followed by bird V, and humans and horses. We assumed that infected mosquitoes do not recover from the infection and that they are not affected by an additional disease-induced death rate. The mosquito population was in equilibrium as the birth rate and the death rate *b_M_* were considered equal.

The bird, human, and horse populations were subdivided into susceptible (respectively *B_VS_*, *B_HS_*, *H_S_* and *E_S_*), exposed (respectively *B_VE_*, *B_HE_*, *H_E_* and *E_E_*), infectious (respectively *B_VI_, B_HI_, H_I_ and E_I_),* and recovered (respectively *B_VR_*, *B_HR_*, *H_R_* and *E_R_*). The bird and horse populations were in equilibrium in the absence of disease as the natural birth rate and the natural death rate (respectively *b_B_*=*m_B_* and *b_E_*=*m_E_*) were considered equal. They were affected by an additional disease-induced death rate *ν_VA_*, *ν_HB,_* and *ν_E,_* respectively, and the count of dead birds V over time is reported in a compartment *B_VD_*. Additionally, WNV was maintained in the population via stochastic events of importation of infectious birds at a rate *Λ_B_*. We considered that the human population dynamics are insignificantly slow compared to the disease dynamics, and we did not include them in the equations. The transmission rate by infectious mosquitoes was respectively *p_M_δ_M_kp_BV_*, *p_M_δ_M_kp_BH_*, *p_M_δ_M_kp_H_*, and *p_M_δ_M_kp_E_* (with *p_M_* the mosquito-to-host transmission probability, *δ_M_* the fraction of non-diapausing mosquitoes, *k* the biting rate of mosquitoes, and *p_BV_*, *p_BH_*, *p_H_* and *p_E_* mosquito biting preference coefficients).

We used the parameter values provided in (28) and (30) as default values (Supplementary Material, section Model details). Intrinsic stochasticity was added to the model using a branching process implemented using the Gillespie algorithm (31). Simulations were run in R 4.2.3 using the package SimInf (32).

### Perturbation-recovery experiments

To study the presence of critical slowing down in this system, we performed perturbation-recovery experiments using the WNV model described above for different values of R_0_ (i.e. the basic reproduction number, defined as the number of secondary cases expected from an infectious individual in a fully susceptible population). R_0_ was calculated using the Next Generation Matrix (NGM) (Supplementary material, section Model details) (21), and the biting rate values were chosen to obtain the desired R_0_ values in the simulations described below. Five infectious individuals were introduced in the disease-free population at *t=5*, and the recovery time to the disease-free equilibrium (i.e. the absence of infectious individuals) was calculated based on the exponential decay of infectious individuals after the perturbation. The slope of a linear regression between the log number of infectious individuals and time with a fixed intercept was used to calculate the decay rate. The reciprocal of this number is the recovery time from a perturbation. We performed these experiments for both birds and mosquitoes.

### Data simulations

For our analyses, we assumed that mosquito incidence, bird incidence and seroprevalence, human incidence and seroprevalence, and horse incidence and seroprevalence could be monitored, initially with high precision.

The biting rate varied over time to alter the R_0_ and, thus, the resilience of the system. The biting rate is known to be influenced by temperature, which mimics a simplified representation of the emergence of WNV due to increasingly suitable climatic conditions (Paz & Semenza, 2013). Other parameters affected by the temperature, such as the extrinsic incubation period or mosquito birth and death rates, are kept fixed to avoid interactions between the parameters obscuring the dynamics of emergence, and thus our understanding of the indicators. For the same reasons, seasonal fluctuations were omitted.

We simulated an approaching epidemic due to increasingly suitable conditions in single long runs, with the biting rate increasing slowly over time. These so-called “emergence time series” were used to measure the performance of the indicators as an early warning for upcoming epidemics. In these time series, R_0_ gradually increased from 0.7 to 1 over 428 weeks with a weekly resolution (Fig 1B). As a control, we simulated “stable time series” under the assumption of a fixed R_0_ over time, leading to no major outbreaks (Fig 1C). We used R_0_=0.8 for that scenario. We simulated 100 stochastic repetitions for each R_0_ condition.

### Monitoring scenarios

We considered 10 data sources to be used in our early-warning signals: mosquito, bird species V, bird species H, human and horse incidence time series represented by model states M_I_, B_VI_, B_HI_, H_I_ and E_I_, bird, human and horse seroprevalence time series represented by the states B_VR_, B_HR_, H_R_ and E_R_ as well as time series of dead birds species V represented by the state *B_VD_*. We considered these time series individually to evaluate the performance of univariate resilience indicators. For multivariate indicators, three scenarios representing different data combinations were created. Scenario A represents an anthro-equine scenario, where all human and horse data (incidence and seroprevalence) are being collected (H_I_, E_I_, H_R_, E_R_). Scenario B represents a wildlife scenario where all mosquito and visible (i.e. detectable) bird data (incidence, seroprevalence and deaths) are being collected (M_I_, B_VI_, B_VR_, B_VD_). Finally, scenario C represents a hidden bird species investigation where the hidden bird species data is collected along with mosquito data (M_I_, B_HI_, B_HR_).

To try to identify what affects the prediction performance of a given species, we vary the feeding preference coefficient, affecting how much each species gets infected by a typical infectious individual. We generated time series of the model with six values for the feeding preference coefficient of mosquitoes towards bird species H. The feeding preference coefficient towards bird H is chosen as it has a direct, large effect on how much all the species get bitten by mosquitoes, and thus can get infected (Supplementary material, Fig S12). Additionally, it allows to explore how potential hidden reservoirs could obscure early-warning signals. For each feeding preference scenario, we quantified how much each species gets infected by a typical infectious individual, called the *typical infection coefficient* in the rest of this manuscript (Fig S12). To estimate the *typical infection coefficient,* we used the corresponding coefficient in the eigenvector associated with the dominant eigenvalue of the NGM (Supplementary material, Section model details). In the default emergence time series, bird H is preferred by the mosquitoes with a coefficient p_BH_=10, against p_BV_=5 for bird V, and p_E_=p_H_=1 for horses and humans. Additionally, we generated time series with p_BH_=5, 20, 30, 50, and 80. For each simulation, we adapted the range of values for the biting rate to keep a range of R_0_ going from 0.7 to 1 for the emergence time series and R_0_=0.8 for the stable time series.

To investigate the limitations of resilience indicators as an early warning sign for upcoming epidemics in data-poor scenarios, we explored the effect of two additional complexities: (a) reduced resolution, and (b) reduced observation probability. For (a), the data were down-sampled from the full emergence time series by reducing the resolution. Thus, the R_0_ gradient remains unchanged, but the number of data points decreases. Secondly, (b) was explored using the full emergence time series. For each data point, observation noise was added using a Poisson distribution of expectation *λ=number of observations* for that time step for that variable. To reproduce imperfect observations, we defined three observation probabilities: *p_1_* = 0.1, *p_2_*=0.01, and p_3_=0.001. Imperfect observations were then generated for each data point using a Poisson distribution of expectation *λ=p_i_ x number of observations*.

### Resilience indicators

After detrending the emergence time series using a Gaussian kernel with the optimal bandwidth according to (Bowman & Azzalini 1997), we analyzed the residuals to calculate the indicators of resilience over time. A rolling window of 50% of the size of the time series was used to assess the trends in the indicators over time (36). We then evaluated the strength of the trend in the indicators over time using the Kendall tau correlation. We used an implementation of the indicators in Matlab (37).

We used a subset of the multivariate indicators described in Weinans et al. (2021). These indicators rely on different techniques to combine multiple time series. Some indicators use the maximum or average of a univariate indicator among all time series. For instance, the maximum variance (Max Var) and maximum autocorrelation (Max AR) calculate the variance (resp. autocorrelation) for all variables but keep only the maximum value of variance (resp. autocorrelation) for each window. Similarly, the mean variance (mean var) and mean autocorrelation (mean AR) take the mean of variance (resp. autocorrelation) over all variables for each window. The maximum covariance (max cov) is the maximum coefficient of the covariance matrix, and the mean cross-correlation (max abscorr) is the mean of all the absolute correlations between the variables. Furthermore, two more elaborate data reduction techniques were used to combine all the time series (PCA and MAF). PCA indicators were calculated in time series of the system projected in the direction of highest variance, corresponding to the first component of a principal component analysis. The variance (PCA var) and autocorrelation (degenerate fingerprinting, Deg Finger) were then calculated in the projected time series. Similarly, the direction of highest autocorrelation was identified using an MAF analysis (Max Autocorrelation Factor analysis, see Supporting Information) (38). The minimum eigenvalue of the MAF analysis (MAF eig) was calculated for each window, and variance (MAF var) and autocorrelation (MAF AR) were calculated for the projected time series. We compared the performance of these multivariate indicators with the most common univariate indicators, namely autocorrelation and variance of each single time series.

We quantified the indicators’ performance as early-warning signals using ROC (receiver operating characteristic) curves. Emergence time series with both R_0_ conditions (increasing R_0_ leading to an epidemic and fixed R_0_ with no upcoming epidemic) were used to calculate the true positive and true negative rate for different cut-off values of the Kendall tau. The Area Under the ROC Curve (AUC) was used to estimate the prediction performance of the different indicators under different data scenarios.

## Results

### Perturbation recovery experiments

To investigate the presence of critical slowing down in the model, perturbation recovery experiments (Fig 2A) were performed by introducing infectious birds (Fig 2A, 2B) and mosquitoes (Fig 2C) to disturb the disease-free equilibrium. In both cases, the recovery time increased as R_0_ approached the critical threshold, indicating a loss of resilience as the system approaches suitable conditions for an epidemic to spark.

**Fig 2.**
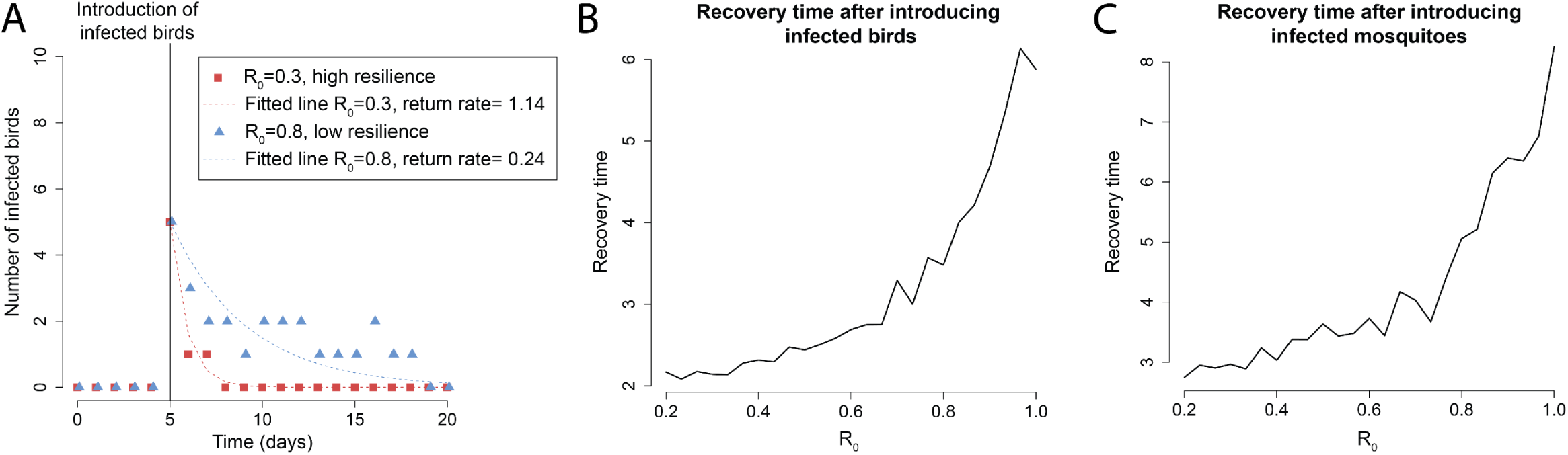
(A) Number of infected birds A over time, for two examples of perturbation-recovery experiments in a case of high (red) and low (blue) resilience. The points represent the observations of infected birds over time, and the dotted lines indicate the fitted lines used to calculate the return rate to the disease-free state (see Methods). (B) Recovery time to the disease-free state after perturbing the system by introducing infected birds for different values of R_0_. (C) Recovery time to the disease-free state after perturbing the system by introducing infected mosquitoes for different values of R_0_. For B and C, the recovery time is defined as the reciprocal of the return rate, assuming exponential decay.

### Performance of the indicators as an early warning for upcoming epidemics

The Area Under the ROC Curve was used as a measure of the performance of the different indicators and scenarios, as it considers both the true positive rate (ability to correctly predict an upcoming epidemic) and the true negative rate (ability to predict that no upcoming epidemic is approaching). For univariate time series, we measured the performance for each time series and univariate indicator (Fig 3A). For multivariate time series, we measured the prediction performance for each monitoring scenario and multivariate indicator (Fig 3B, 3C and 3D). The performance of univariate indicators ranges between 0.68 (Horse seroprevalence with autocorrelation) and 0.88 (Human incidence with variance). The performance of multivariate indicators ranges between 0.56 (Wildlife scenario with explained variance) and 0.94 (Hidden reservoir scenario with mean variance). Although the Hidden reservoir scenario yielded the best prediction performance across all indicators, there were no meaningful differences between the average prediction performance of the different multivariate scenarios. Interestingly, the difference in average prediction performance between the univariate indicators and the multivariate ones was only 0.02 (AUC of 0.79 for univariate indicators and 0.81 for multivariate, average across all scenarios and indicators), especially considering that multivariate indicators use up to four times more data points than the univariate ones, which requires a greater monitoring effort.

**Fig 3.**
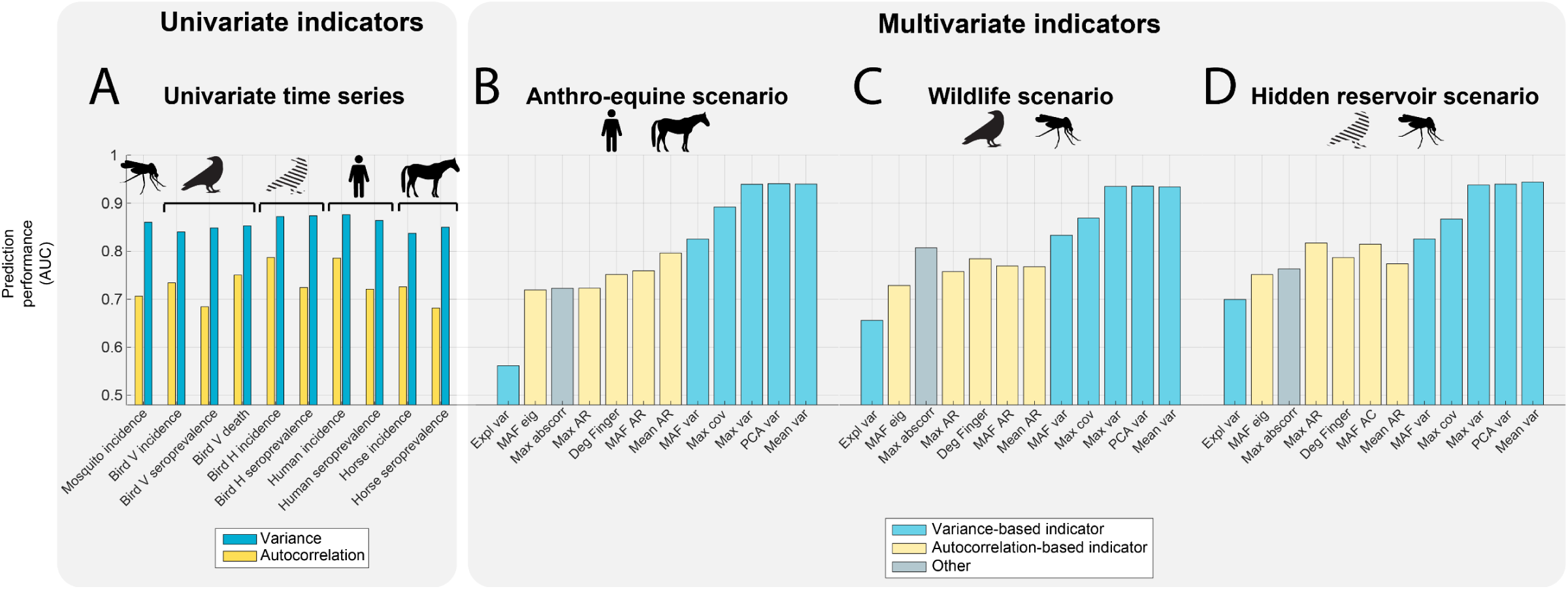
Prediction performance of the different univariate and multivariate indicators of resilience evaluated using the AUC. (A) Prediction performance of the univariate indicators for all univariate time series. (B) Prediction performance of the multivariate indicators for the Anthro-equine scenario. (C) Prediction performance of the multivariate indicators for the wildlife scenario. (D) Prediction performance of the multivariate indicators for the hidden reservoir scenario.

Variance and variance-based indicators outperformed autocorrelation and autocorrelation-based indicators for both univariate and multivariate monitoring scenarios. For univariate time series, variance yielded an average AUC of 0.86 over all the time series, against an average AUC of 0.73 for autocorrelation. Similarly, for multivariate indicators, variance-based indicators yielded an average AUC of 0.86 against an average AUC of 0.76 for autocorrelation-based indicators, and the three best-performing multivariate indicators were variance-based (mean variance, PCA variance and max variance). Due to the scarcity of cases far from the epidemic threshold, the time series contained long stretches of consecutive zeros, resulting in a highly variable autocorrelation, which altered the performance of autocorrelation and autocorrelation-based indicators (Fig S1 to S3).

For the rest of the analyses, we proceeded with the three best-performing indicators (variance for univariate indicators, and mean variance, PCA variance and maximum variance for multivariate indicators), the best multivariate scenario (hidden reservoir), and the best univariate time series (mosquito incidence, bird H incidence, seroprevalence, and humans prevalence). The results of the other indicators and scenarios are provided in the Supplementary Material.

### Data-poor scenarios

The performance of univariate indicators of resilience was compared to the performance of multivariate indicators of resilience in down-sampled time series and for lower reporting probabilities. The comparison was performed for different numbers of data points obtained by subsampling the original time series by reducing the resolution (Fig 4A and 4C, other scenarios and indicators in Supplementary Material, Fig S4 to S7) and for different observation probabilities implemented using a Poisson distribution (Fig 4B and 4D, other scenarios and indicators in Supplementary Material, Fig S8 to S11). The prediction performance of all indicators and monitoring scenarios decreased for lower numbers of data points and lower observation probabilities. The performance of multivariate indicators remained higher than that of univariate indicators, and the difference in prediction performance increased when both the resolution and observation probability were further reduced. Autocorrelation-based indicators were more sensitive to a reduced resolution, consistent with previous findings (36) (Supplementary Material).

**Fig 4.**
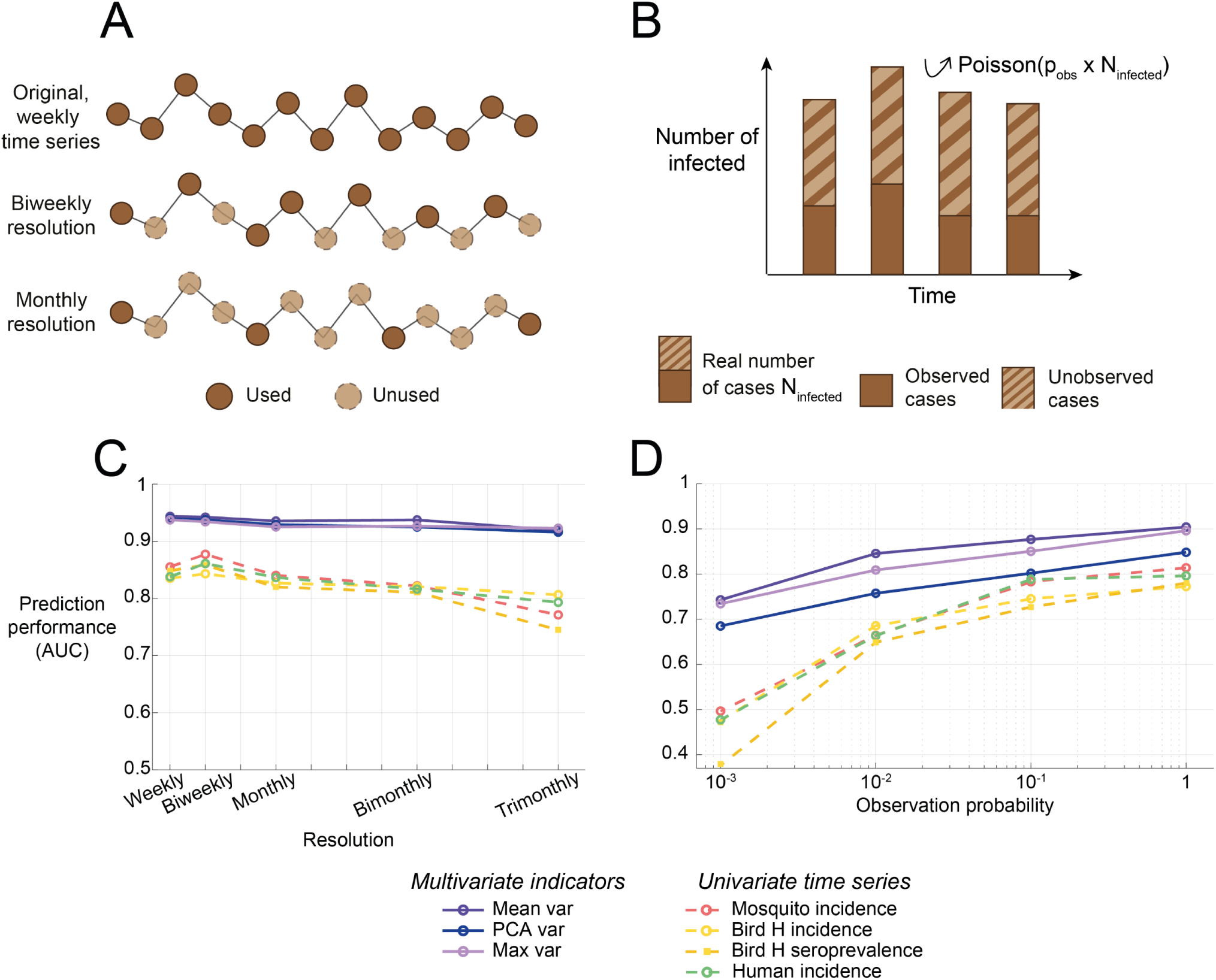
Performance of the best-performing, variance-based indicators of resilience in data-poor scenarios. (A) The resolution is reduced by downsampling the original time series. (B) The observation probability is reduced by subsampling the number of infected using a Poisson distribution. (C) Effect of the reduction of resolution on the prediction performance. (D) Effect of the reduction of observation probability on the prediction performance.

### Variability in the prediction performance of the different species

We explored the nature of the variations in the prediction performance of the different monitoring scenarios observed in the previous section. Specifically, we explored how informative the different species are for early warnings depending on how much they get infected in the transmission cycle. A variation in the mosquito feeding preference towards bird H impacts the *typical infection coefficient* of all the species, i.e. how much they get infected, quantified using the corresponding value in the eigenvector associated with the dominant eigenvalue of the NGM (Fig S12). When the feeding preference towards bird H increased, the prediction performance slightly decreased for univariate indicators uniformly for all species. However, it remained stable for variance-based multivariate indicators (Fig 5A, other scenarios and indicators in Supplementary Material, Fig S13 to S16). Additionally, the prediction performance of the different univariate time series was unrelated to the *typical infection coefficient* (Fig 5B, other scenarios and indicators in Supplementary Material, Fig S17). These results were robust to a reduced observation probability and resolution (Supplementary Material, Fig S18 and S19). Additionally, to make sure that the choice of the varying parameter does not impact the result, the same analyses were repeated by varying the relative abundances of bird H to bird V instead of the feeding preference. These analyses yielded analogous results (Supplementary Material, Fig S20 to S25).

**Fig 5.**
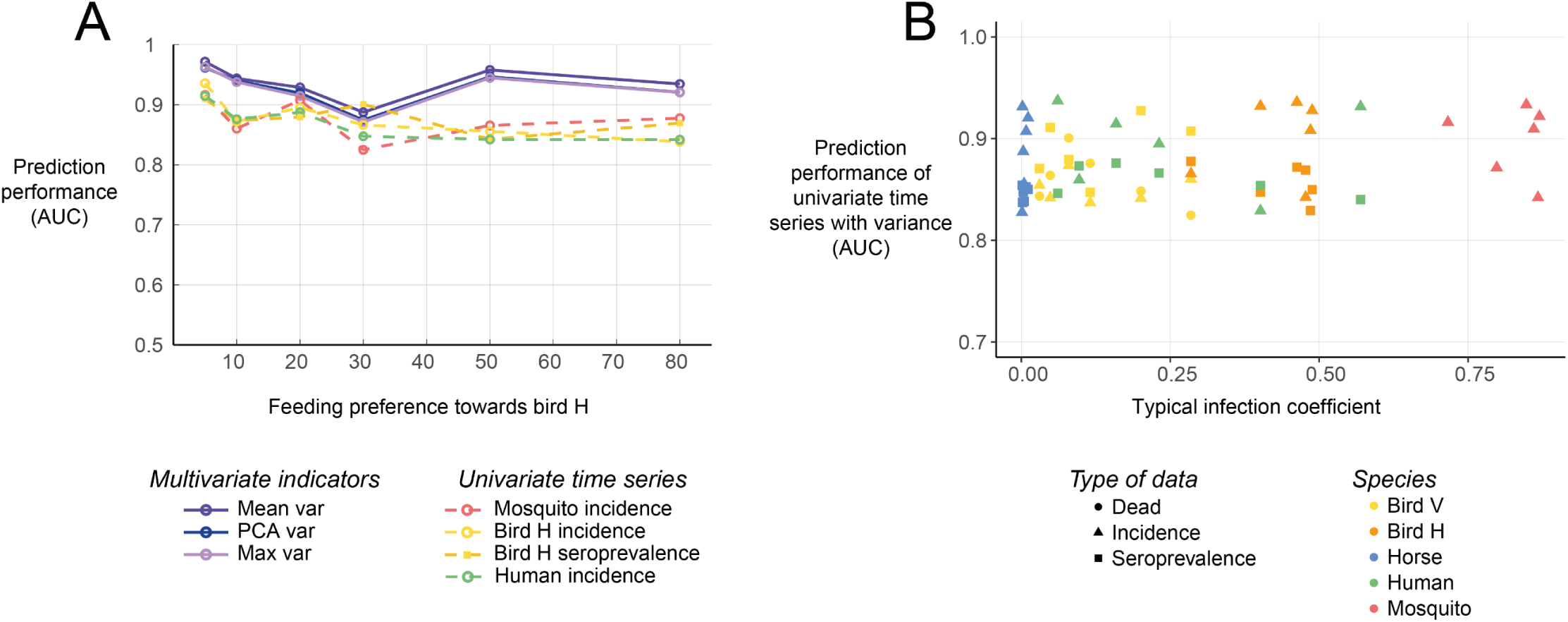
Prediction performance of the different species depending on how much they get infected. (A) Effect of varying mosquito feeding preferences towards the hidden reservoir on the prediction performance. (B) Prediction performance of the univariate time series depending on how much the species get infected, quantified using the *typical infection coefficient* (i.e. the corresponding coefficient of the eigen vector associated with the dominant eigenvalue of the NGM).

## Discussion

Using a synthetic dataset, we found that both univariate and multivariate resilience indicators could signal upcoming outbreaks of WNV. Multivariate indicators outperformed univariate indicators, especially in data-poor scenarios. The accuracy of multivariate indicators ranged up to 0.94, and those of univariate indicators up to 0.88, assuming perfect observation (Fig 3). When reducing the resolution or the observation probability, the difference in prediction performance between univariate and multivariate indicators broadened, suggesting that multivariate indicators are more robust to a decrease in the data quality (Fig 4). However, multivariate indicators require up to four times more data points as several hosts have to be monitored simultaneously. This study was the first to compare multivariate and univariate indicators, and the findings could have important implications for the surveillance of vector-borne diseases. As the model was kept purposefully generic, these results can be extended to other vector-borne diseases. However, future research should confirm the findings in data, including more complexities such as seasonality (39) or complex observation processes (40), as well as cost estimations and logistic ramifications of the different monitoring strategies (21).

The prediction performance of the different multivariate monitoring scenarios, as well as univariate time series, remained stable regardless of how much a species is involved in the transmission cycle. Noticeably, increasing the mosquito feeding preference towards a given species or the relative abundances of the different species did not increase the potential for that species to signal upcoming outbreaks (Fig 5A). Thus, the variability in the prediction performance of the different monitoring scenarios could not be explained by the typical infection coefficient, namely how much a species gets infected by a typical infectious individual (Fig 5B). In contrast to earlier findings, in this system, we found no evidence that species that are central to the transmission cycle are more informative for early warning (14,19). A possible explanation could be the relatively high spillover rate to dead-end species assumed in the model, which is not always observed in practice (39). Additional research is needed to better understand what drives how informative a data source is in anticipating future disease outbreaks using resilience indicators. Answering this question is especially relevant for the surveillance of various infectious diseases, especially when silent transmission is ongoing in hidden reservoirs that are hard to monitor (40).

Variance-based indicators outperformed autocorrelation-based indicators and remained robust in data-poor contexts. This is consistent with previous studies that identified variance or related indicators as the best-performing indicators when researching the performance of resilience indicators to anticipate epidemics (5,41,42). In the present study, the poor performance of autocorrelation-based indicators was likely due to the excess of zeros, mainly when R_0_ is far from the critical threshold, leading to a high autocorrelation and thus obscuring the overall trend in autocorrelation (Fig S1 to S3). Additionally, the performance of autocorrelation-based indicators decreased when the resolution of the time series was reduced, in agreement with previous studies (11,43) (Fig S4 to S6). Multivariate variance-based indicators of resilience showed stable performance, even when reducing the resolution of time series or the observation probability (Fig 4). This was especially true for indicators using the average of univariate indicators, consistent with previous research (11). Contrastingly, explained variance performed particularly poorly, probably because of the reactivity of the system (14). As data requirements are a major factor limiting the use of resilience indicators in practice, multivariate indicators hold potential to improve the performance of resilience indicators as an early warning of upcoming epidemics using fewer data.

This study was limited to synthetic data. The model was purposefully kept simple to ensure the genericity of the results. The emergence of the virus was driven by biting rates increasing in response to increasing temperature. However, temperature has an impact on other parameters, such as mosquito population dynamics parameters and the extrinsic incubation period. However, resilience indicators are found to be robust to the choice of bifurcation parameter as long as the change is gradual (43). Further, seasonal fluctuations were omitted. Previous research with univariate indicators found a limited impact of periodicity on the prediction performance of resilience indicators (41), which is likely to hold for multivariate indicators. Although indicators of resilience loss hold potential to improve the current practice of early warning for vector-borne diseases, these findings should be confirmed in datasets generated with complex models, but also real data specific to a certain pathogen.

Finally, multivariate indicators require sampling of many data sources simultaneously. This is complicated by cost and logistic limitations, but also coordination of the various authorities. The different data sources are often monitored by different institutions, such as public health authorities for human cases, veterinary authorities for domestic animals, and environmental authorities for wildlife. Coordinating the monitoring of different authorities and sharing data between the different institutions is sometimes challenging (44). Additionally, costs and logistics limit the monitoring of wild birds and mosquitoes, and it is thus impossible to cover a whole, often heterogeneous area. This was omitted in the model, which assumed homogeneous mixing as well as homogeneous spatial coverage of wildlife data. In the case of spatial disparities in transmission, it is essential to consider high-risk zones when choosing where to execute the monitoring (45). For instance, West Nile transmission can be affected by the local ecological context, such as the availability and abundance of different (competent and non-competent) hosts and vectors (27), leading to spatial heterogeneities in transmission potential (46). Finally, some data sources studied here inherently contain biases that were not accounted for in the model. The model assumed that the number of infected mosquitoes was reported over time. In practice, the number of infected mosquito pools is usually reported, reflecting the presence or absence of infected mosquitoes in a given area and not the number of infected mosquitoes (9). Seroprevalence data is primarily informative about the past state of infection and could introduce biases if not sampled at regular intervals (23,47). Additionally, when multiple similar pathogens are circulating simultaneously, antibodies targeting one pathogen may also react to a different, similar pathogen in tested individuals. This phenomenon, called cross-reactivity, can lead to a non-random overestimation of seroprevalence in areas where several pathogens are co-circulating. The dynamics of the co-circulating pathogen could be captured in the seroprevalence data, potentially introducing biases when measuring resilience (48). Infected mosquito pools and seroprevalence data should thus be used with caution. While multivariate indicators have the potential to solve the challenge of large data requirements by monitoring several data sources instead of focusing on one sole source of information, many considerations remain in designing adapted surveillance programs. The final decision should be based on the specific challenges of a given area and pathogen, as well as whether the additional effort is worth the gain in prediction performance.

## Conclusion

This study was the first to show that multivariate indicators of resilience outperform univariate indicators in anticipating future outbreaks of mosquito-borne diseases, especially in data-poor contexts. As high granularity and quality of required data limit the use of resilience indicators in infectious disease epidemiology in practice, multivariate indicators hold the potential to improve the prediction performance of resilience indicators while dealing with data-poor contexts. However, adapting the surveillance programs to collect the relevant data for multivariate indicators of resilience entails new challenges related to costs, logistic ramifications and coordination of different institutions involved in surveillance. Ultimately, the aim is to inform effective monitoring strategies, and this study represents an initial step towards that objective.

## Funding

This publication is part of the project ‘Preparing for Vector-Borne Virus Outbreaks in a Changing World: a One Health Approach’ (NWA.1160.18.210), which is (partly) financed by the Dutch Research Council (NWO).

## Acknowledgements

We thank Marten Scheffer for the insightful discussions, his help with interpreting the results, and his contribution to the funding acquisition. We thank Els Weinans for providing her implementation of the multivariate indicators of resilience and taking the time to clarify some details about the methodology. We also thank the members of Work Package 3 of the One Health Pact consortium, Mariken de Wit, Afonso Dimas Martins, Mart de Jong, Hans Heesterbeek, and Martha Dellar, for their interest in the project and suggestions to improve this work.

## Supplementary material

### Model details

#### Equations

Model equations are based on a simplified version of the model provided in (Laperriere et al., 2011).

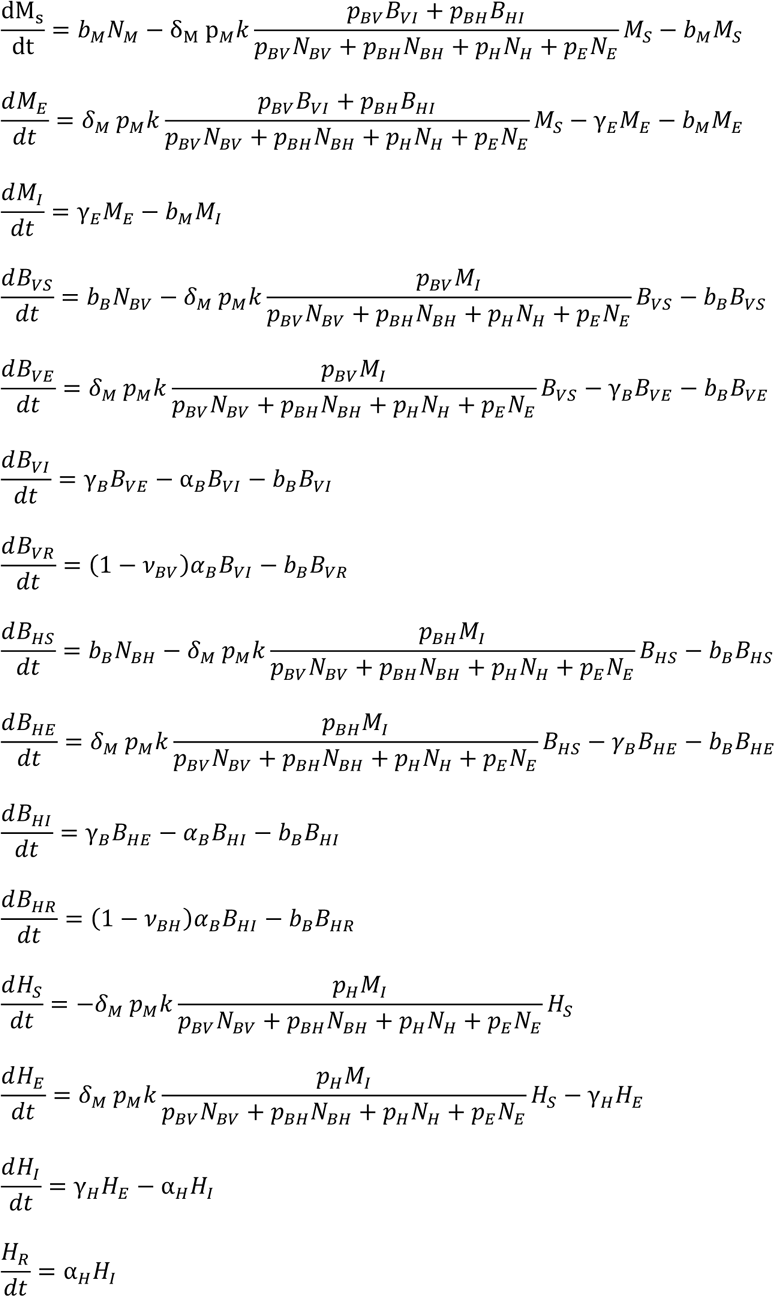

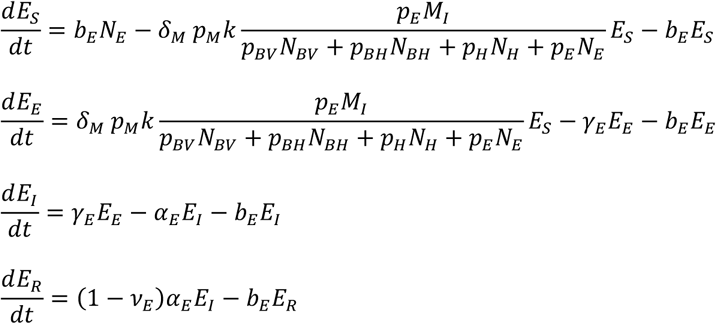

#### Parameter values

**Table S1:**
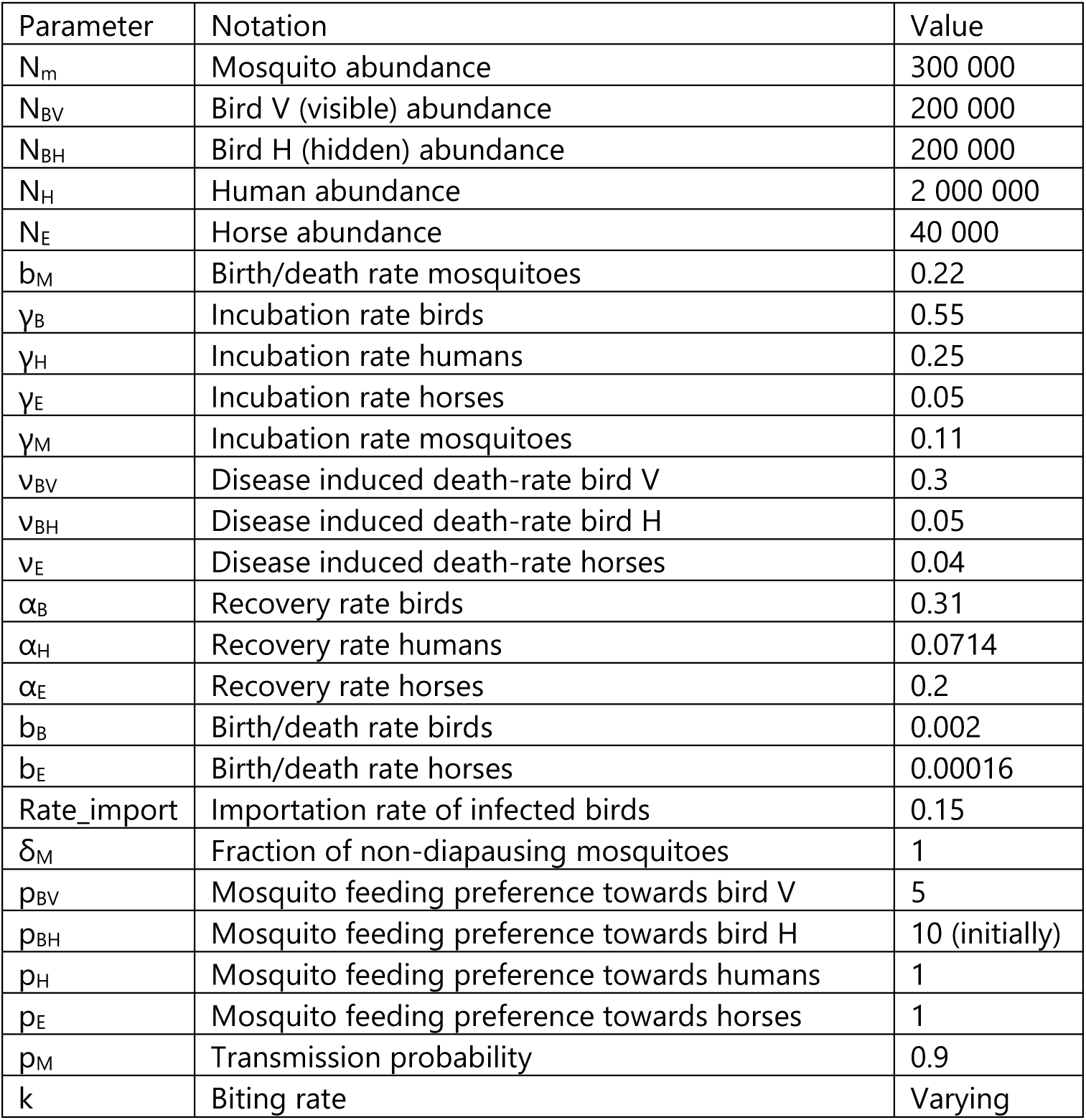
Parameter values for the model. Parameters are taken from (de Wit et al., 2024; Laperriere et al., 2011).

#### Next Generation matrix

The Next Generation matrix was used to estimate R_0_ as well as the typical infection coefficient calculated in the eigenvector associated with the dominant eigenvalue of the NGM. The NGM with large domaine is calculated numerically using the formula provided in (Diekmann et al., 2010): K_L_=-T Σ^-1^, with Σ being the transition matrix and T the transmission matrix. T and Σ are defined as follows for our system:

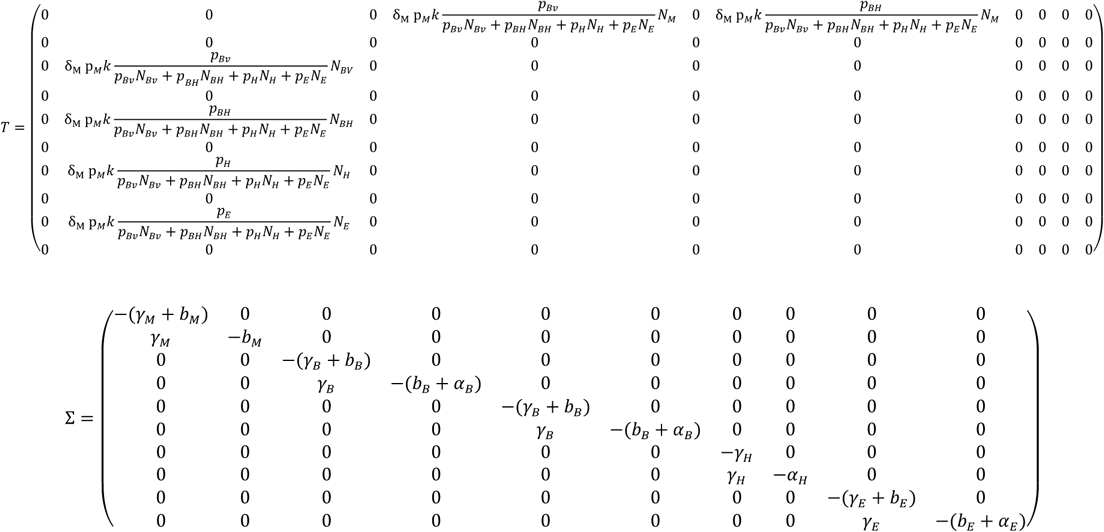

The NGM is obtained by multiplying the NGM with large domain by an auxiliary matrix E, as defined in (Diekmann et al., 2010). The eigenvalues and eigenvectors of the NGM are calculated numerically using R 4.2.3.

### Multivariate Factor Analysis (MAF)

The main essence of Multivariate Factor Analysis (MAF) is to detect the direction of highest autocorrelation, similarly to a PCA with variance (Weinans et al., 2019, 2021).

To find the direction of highest autocorrelation, we proceed as follows:

1. The dataset is transformed to ensure it has an identity matrix as the covariance matrix, using an SDS transform (Haugen et al., 2015). X_SDS_ is the resulting dataset.
2. The first difference of X_SDS_, [X_SDS_(t)-X_SDS_(t+1)], is calculated
3. The eigenvector V and eigenvalues E of the covariance matrix of the first difference [X_SDS_(t)-X_SDS_(t+1)] are calculated
4. The dataset is projected in the direction of highest autocorrelation by multiplying by V

It is important to note that MAF is inverted compared to a PCA: the minimum eigen value of the MAF corresponds to the maximum eigen value for a PCA.

### Multivariate indicators of resilience for different R0

To test the behaviour of our indicators for different values of R_0_, we generated time series under different fixed R_0_ values under the critical threshold (i.e. <1), and analyse each time series separately. We call these data “fixed R_0_ runs”. Each of these time series was simulated over 70 weeks, with a weekly resolution, for 100 values of R_0_ between 0.2 and 1. For each value of R_0_, 100 stochastic repetitions were simulated. The value of the different indicators was then calculated in each repetition for each value of R0, and the median and 95% interval were determined for each value of R_0_. This procedure was repeated for each monitoring scenario and each indicator (univariate and multivariate).

The slowing down observed in the perturbation recovery experiments (Figure 2) was reflected by an increase in resilience indicators in our simulated fixed time series (Figure S1 to S3). Especially variance-based indicators such as mean variance, max variance and PCA variance displayed a strong increase as R_0_ increased. Autocorrelation based indicators such as MAF autocorrelation, mean autocorrelation and max autocorrelation displayed a weaker increase. Additionally, autocorrelation-based indicators sometimes were high for low values of R_0_. Due to the scarcity of cases far from the epidemic threshold, the time series contained long stretches of consecutive zeros, resulting in a highly variable autocorrelation, but these zeros obviously did not reflect any measure of the recovery speed of the system.

**Figure S1.**
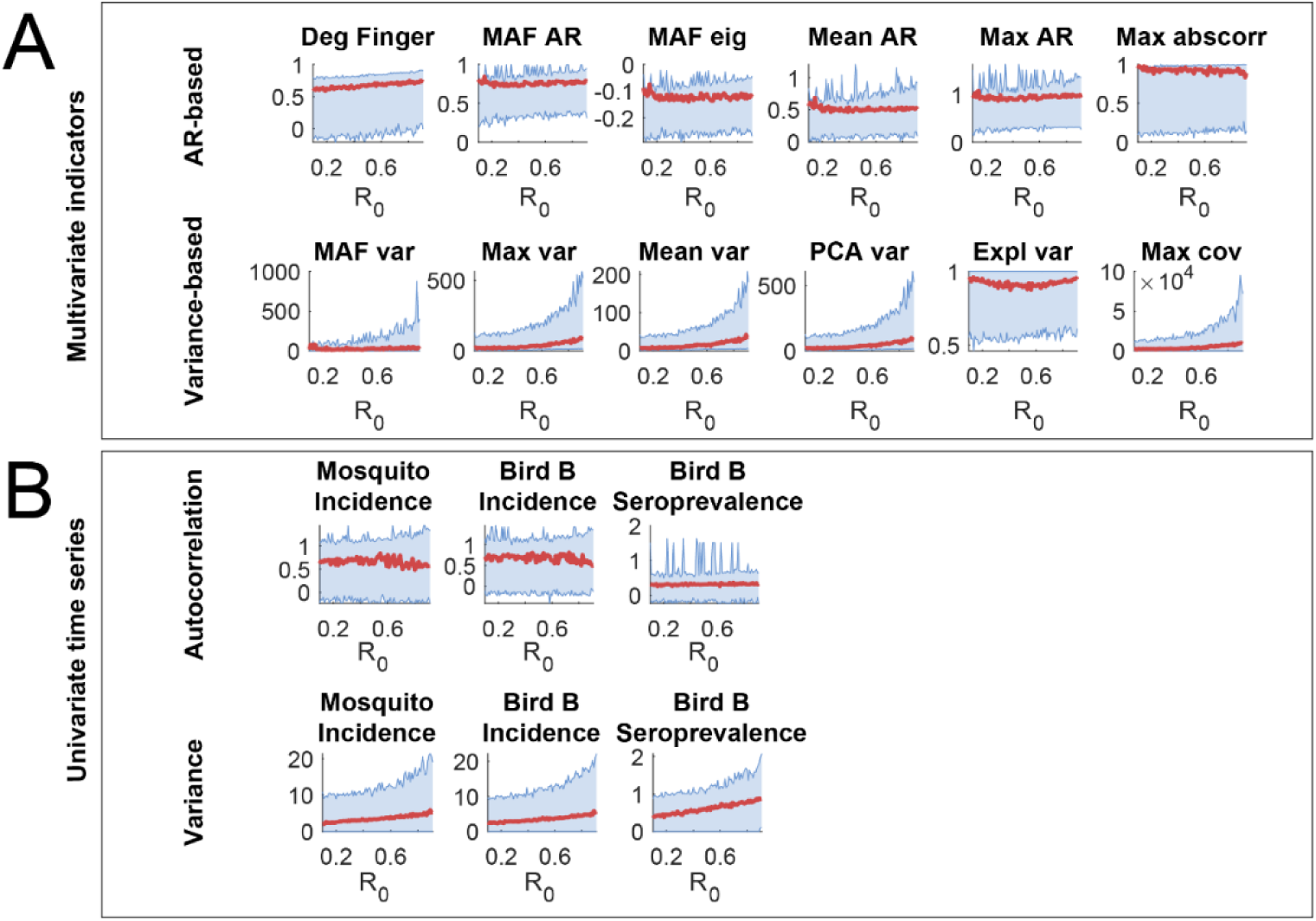
Indicators of resilience, calculated from separate time series generated for each level of R_0_ (i.e. fixed R_0_ runs) following the hidden reservoir monitoring scenario. The red line indicates the median estimation over all the repetitions, and the blue lines are the 2.5 and 97.5 percentiles. (A) Multivariate indicators of resilience, calculated in fixed runs of the multivariate time series following the hidden scenario. (B) Univariate indicators of resilience calculated in fixed runs of univariate time series contained in the hidden reservoir scenario.

**Figure S2.**
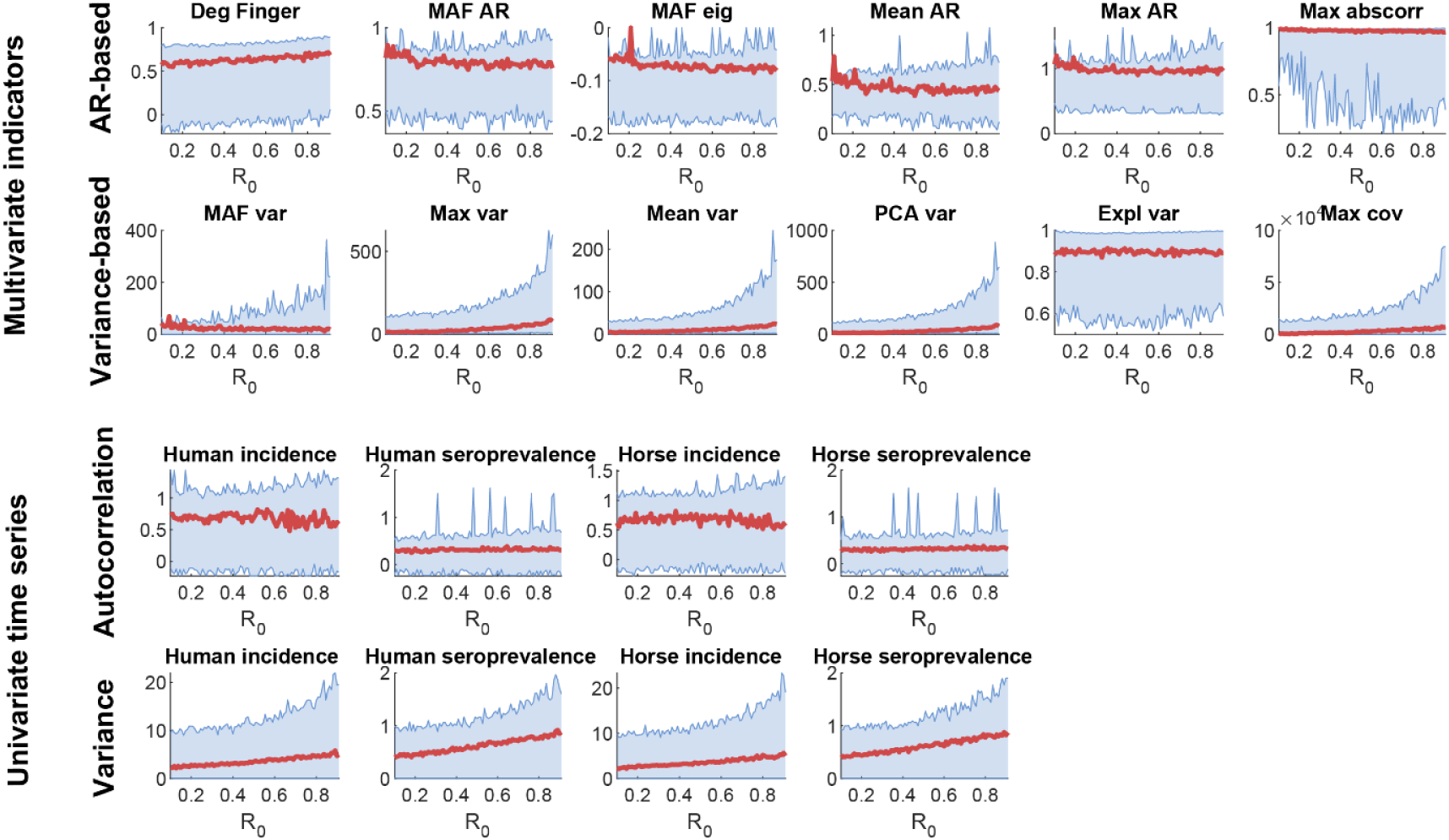
Multivariate indicators of resilience, calculated from separate time series of the anthro-equine scenario generated for each level of R_0_ (i.e. fixed R_0_ runs, see Methods). The solid line indicates the median estimation over all the repetitions, and the dotted lines are the 2.5 and 97.5 percentiles.

**Figure S3.**
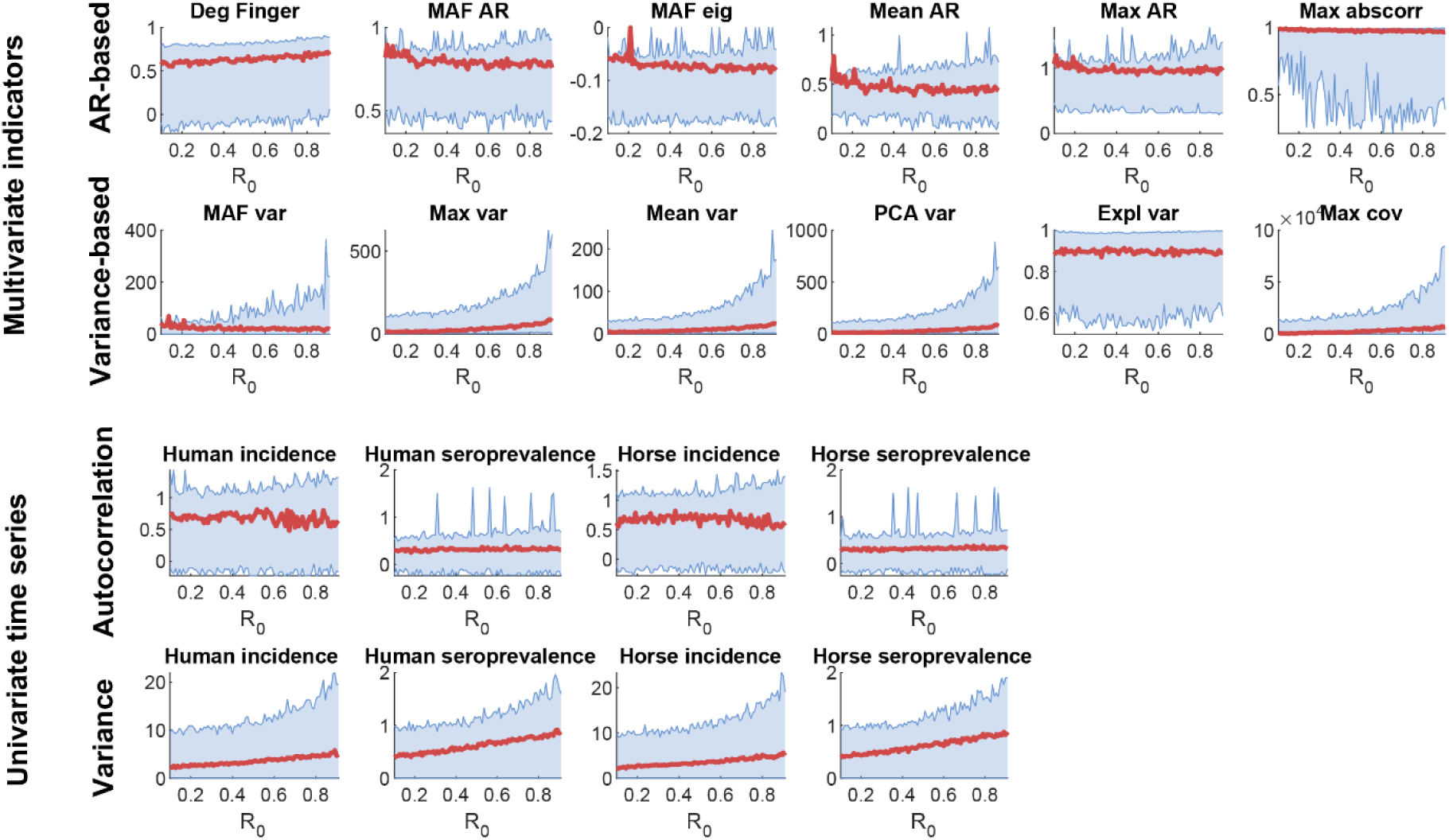
Multivariate indicators of resilience, calculated from separate time series of the wildlife scenario generated for each level of R_0_ (i.e. fixed R_0_ runs, see Methods). The solid line indicates the median estimation over all the repetitions, and the dotted lines are the 2.5 and 97.5 percentiles.

### Down sampling of the data for all the indicators and monitoring scenarios

**Figure S4.**
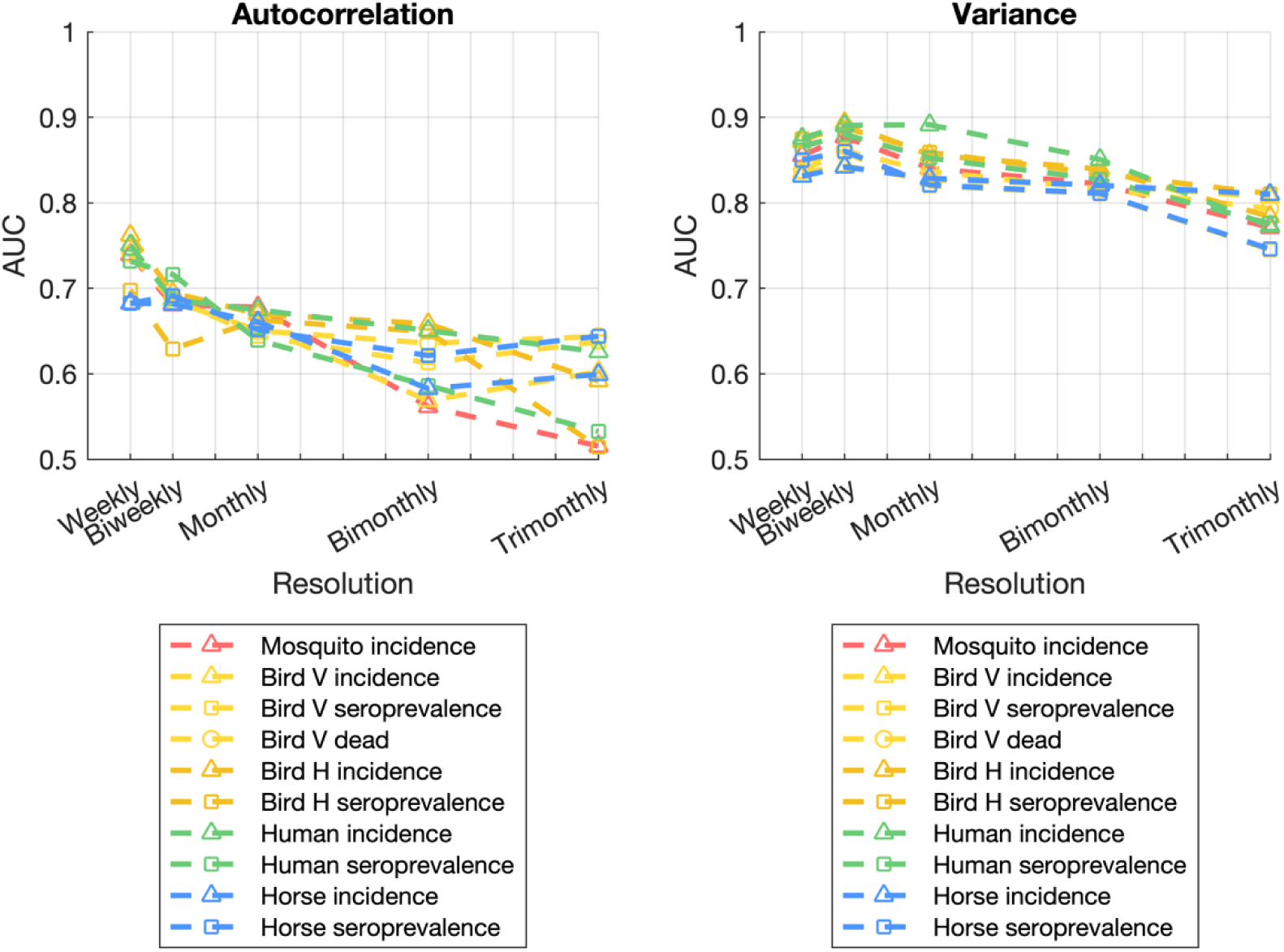
Prediction performance all univariate time series for both autocorrelation and variance, depending on the resolution of the data.

**Figure S5.**
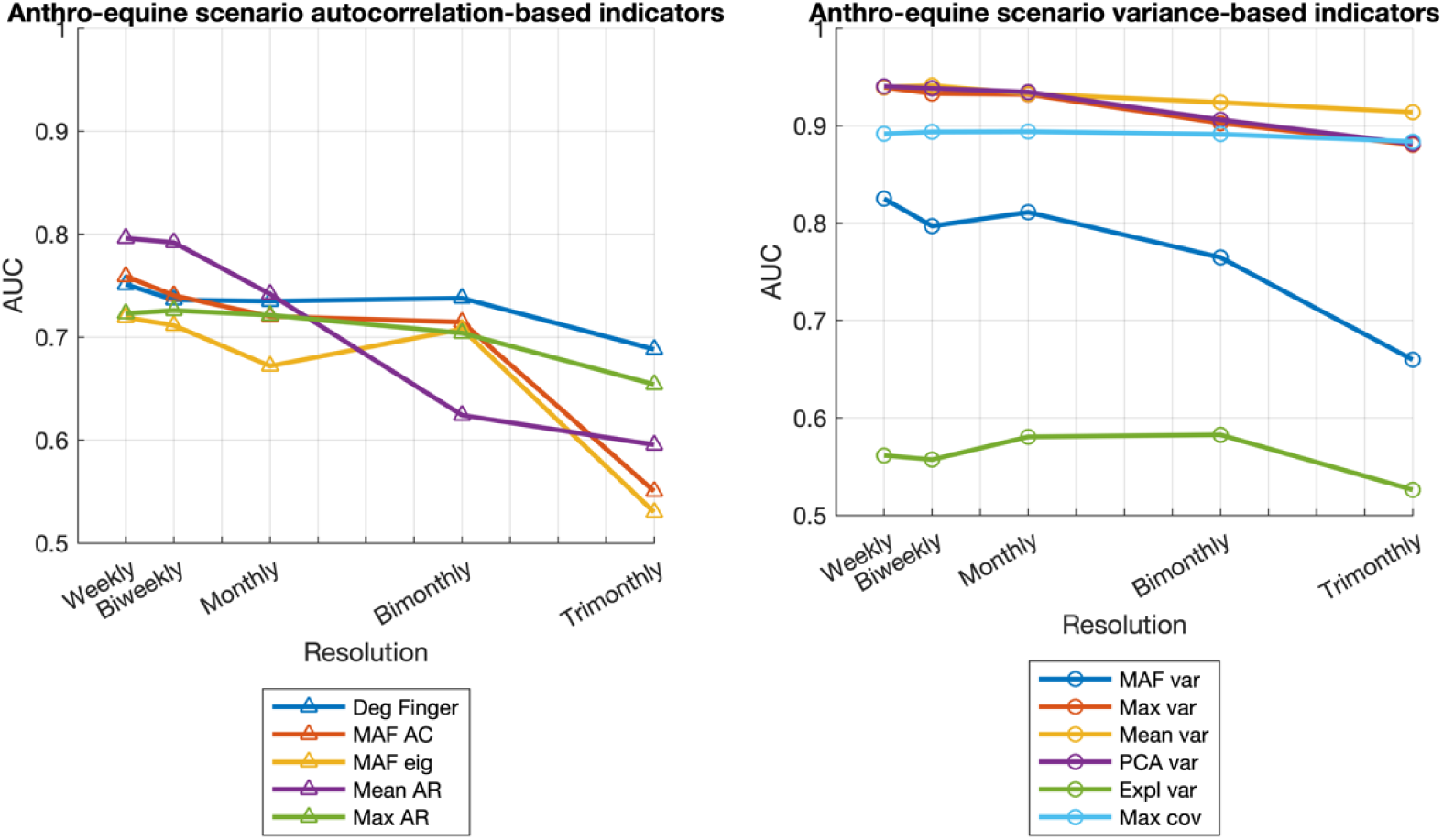
Prediction performance for the anthro-equine scenario for all autocorrelation and variance-based indicators, depending on the resolution of the data.

**Figure S6.**
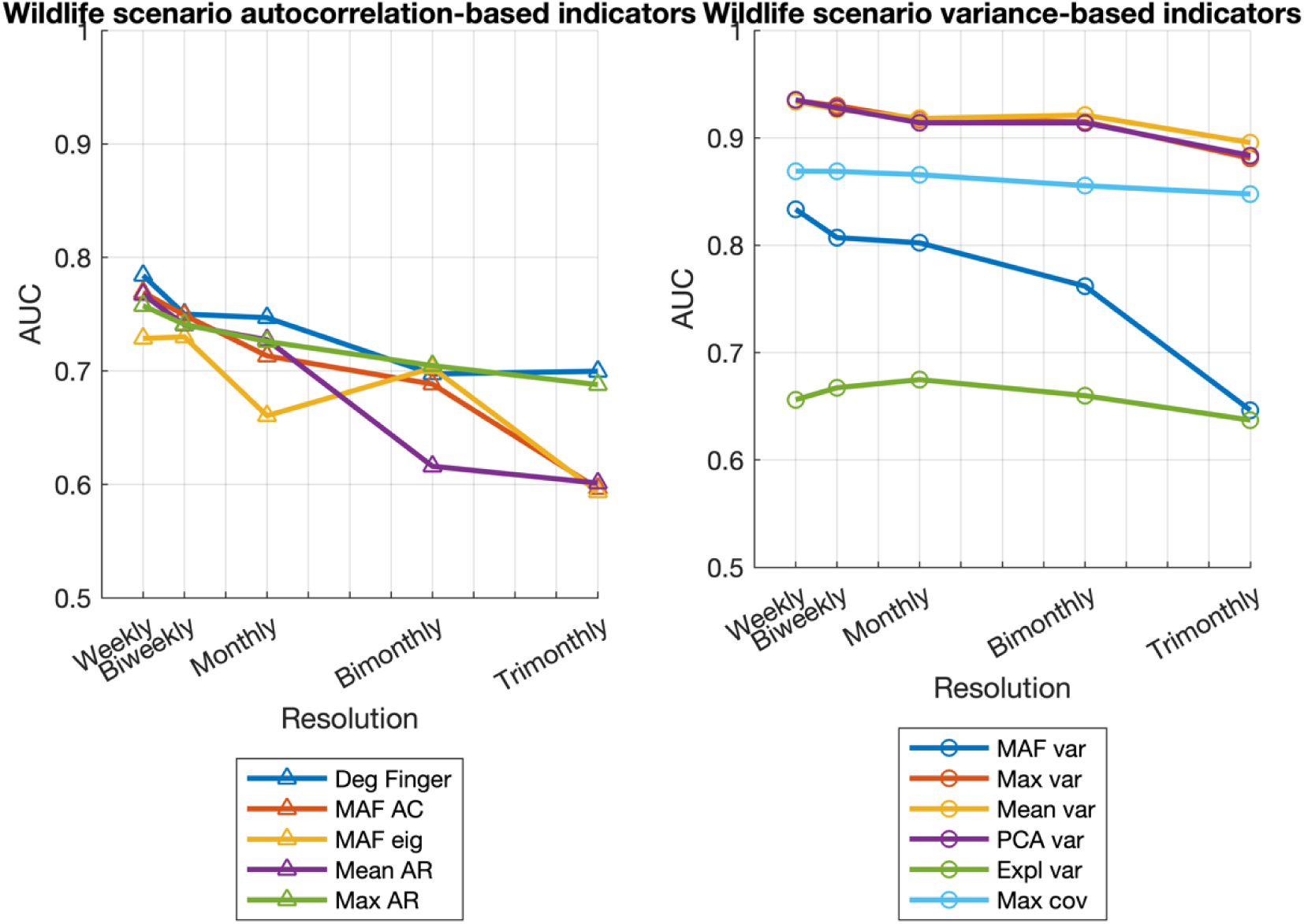
Prediction performance for the wildlife scenario for all autocorrelation and variance-based indicators, depending on the resolution of the data.

**Figure S7.**
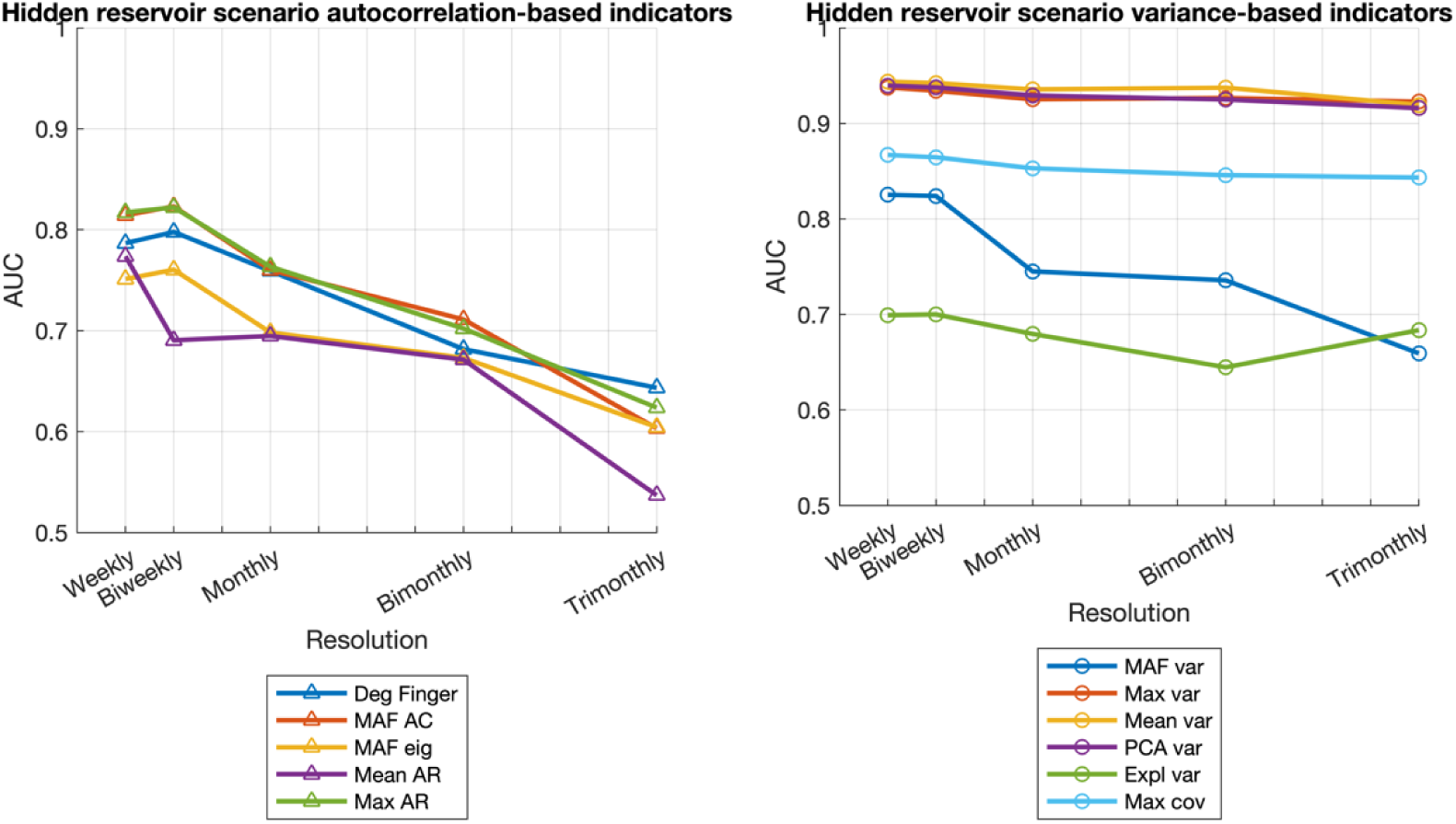
Prediction performance for the hidden reservoir scenario for all autocorrelation and variance-based indicators, depending on the resolution of the data.

### Reducing the observation probability for all the indicators and monitoring scenarios

**Figure S8.**
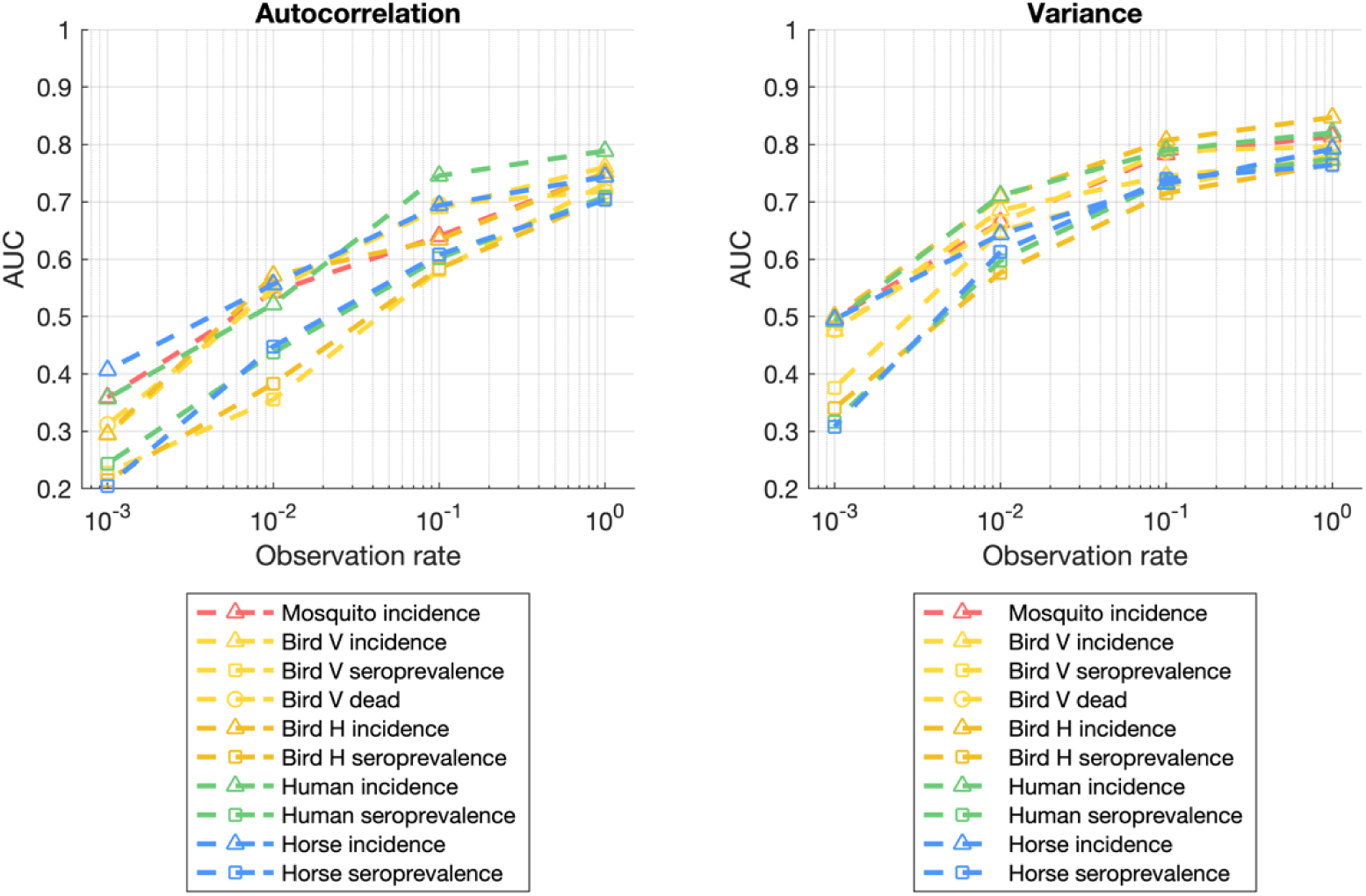
Prediction performance all univariate time series for both autocorrelation and variance, depending on the observation probability.

**Figure S9.**
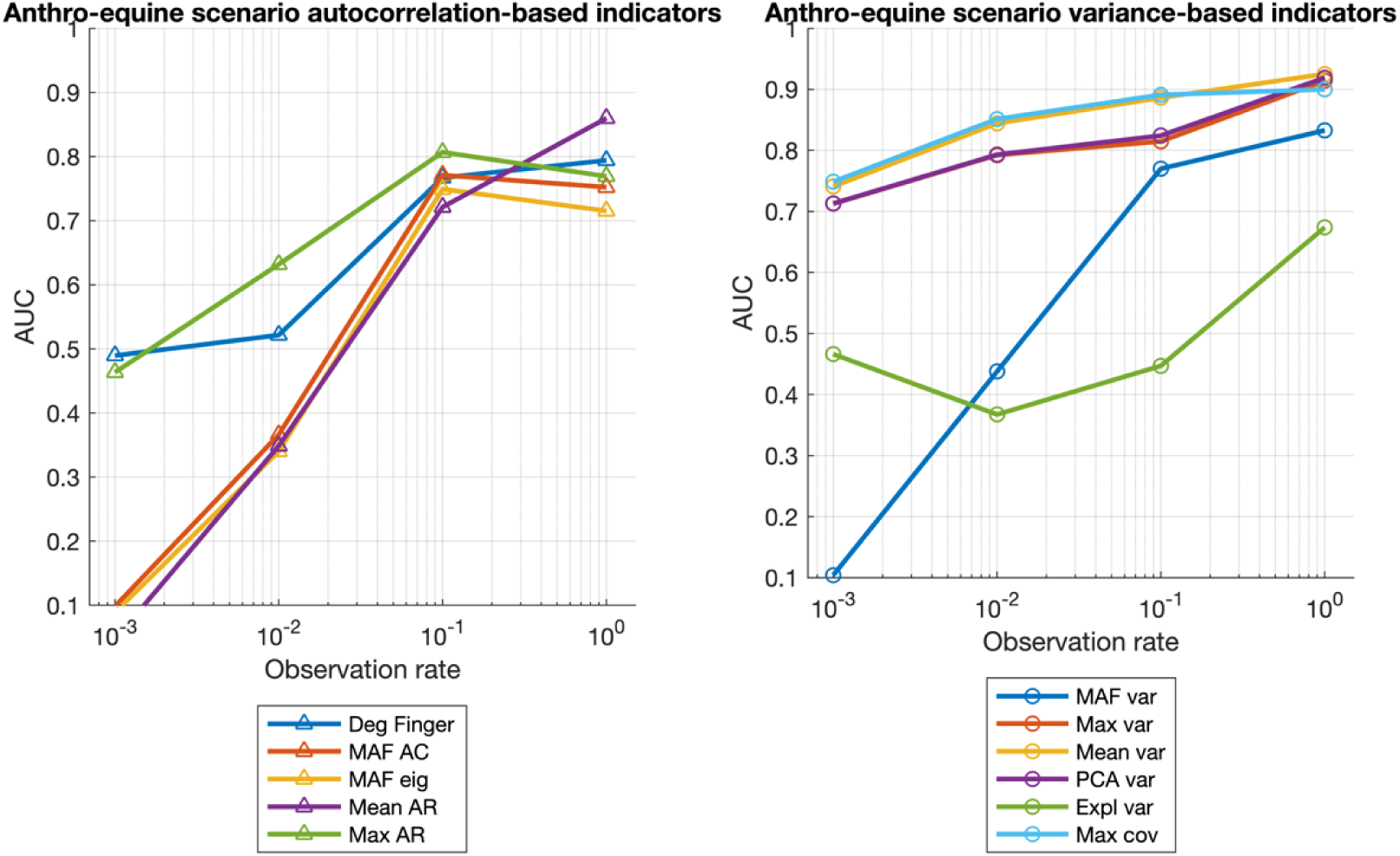
Prediction performance for the anthro-equine scenario for all autocorrelation and variance-based indicators, depending on the observation probability.

**Figure S10.**
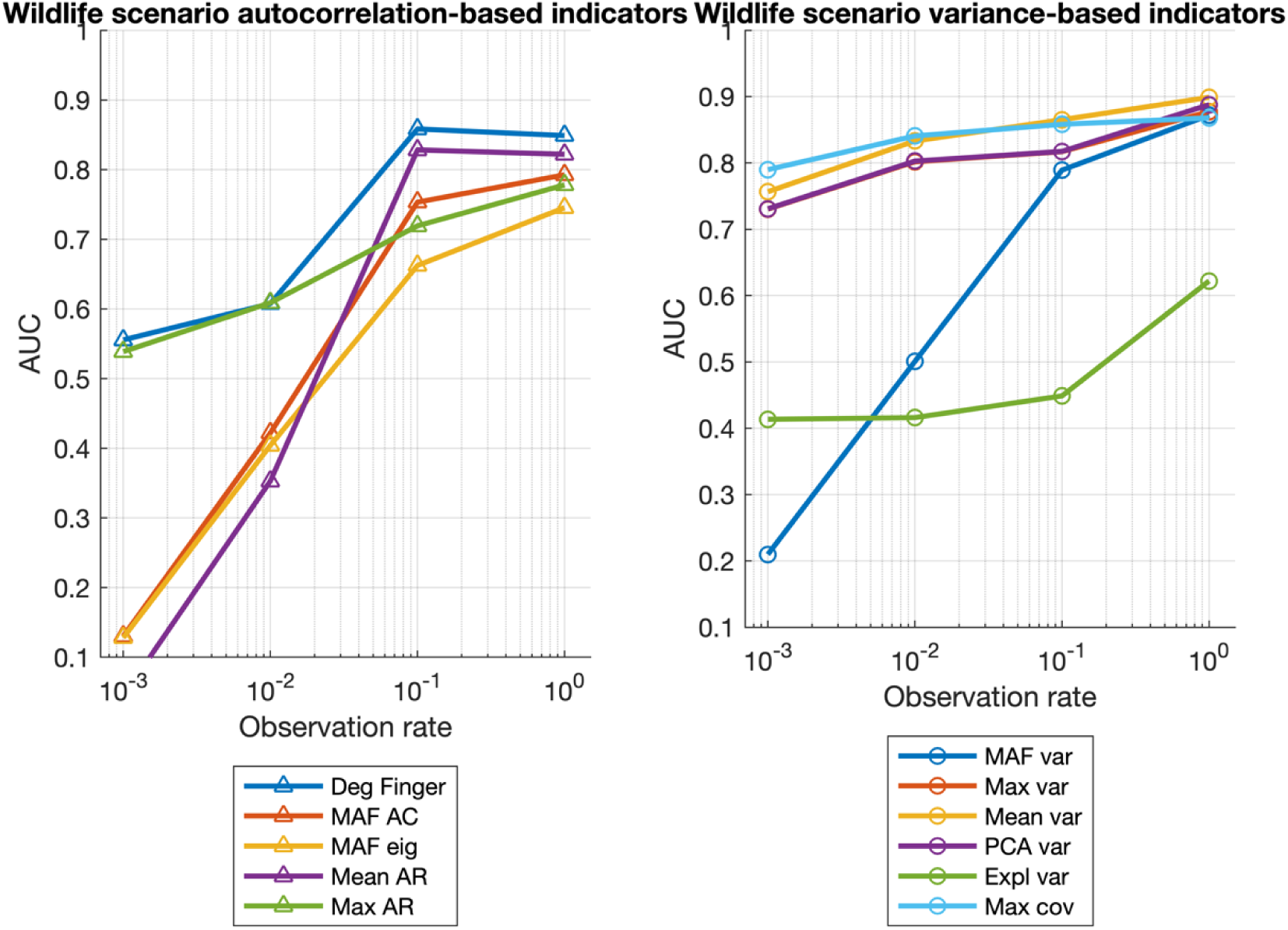
Prediction performance for the wildlife scenario for all autocorrelation and variance-based indicators, depending on the observation probability.

**Figure S11.**
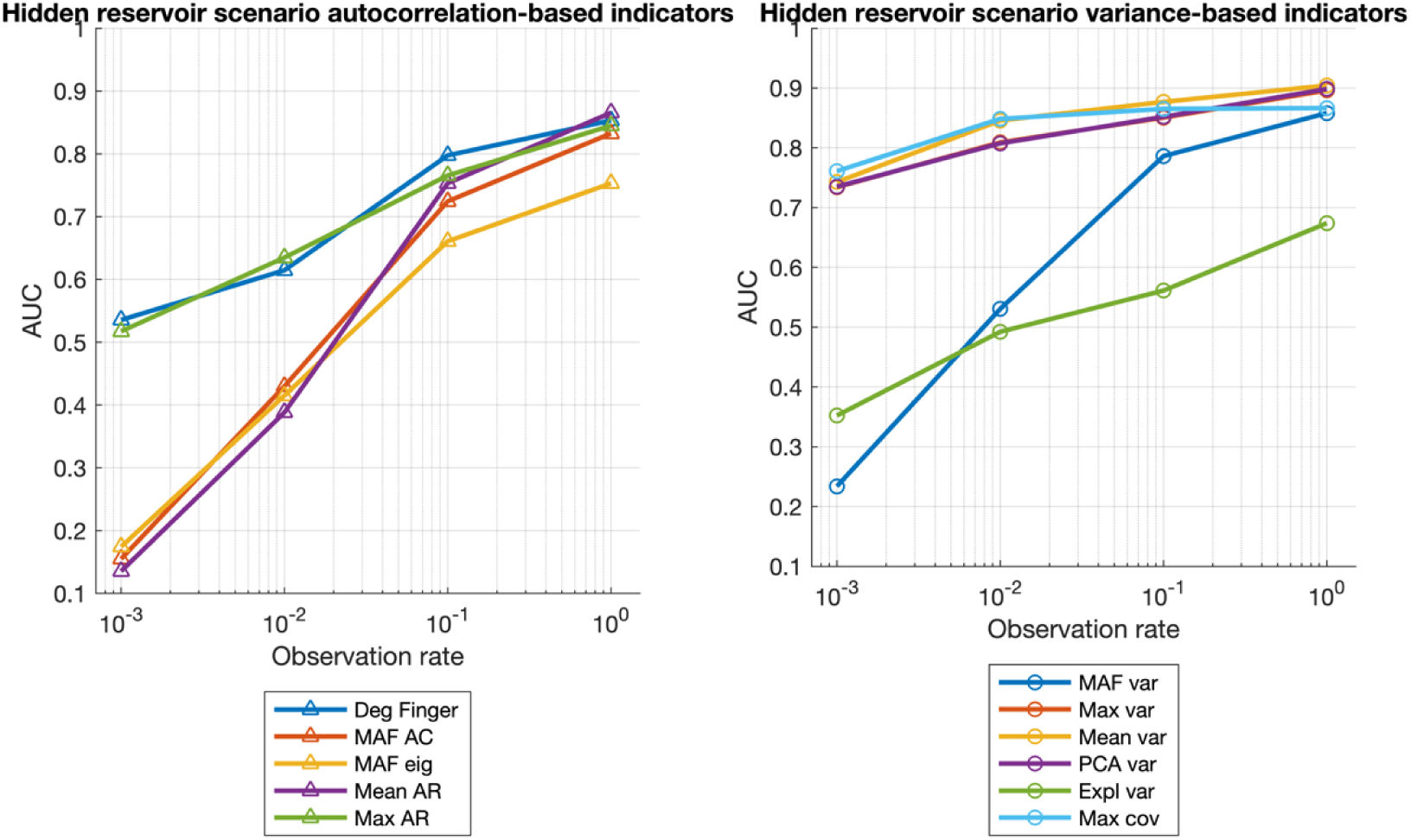
Prediction performance for the hidden reservoir scenario for all autocorrelation and variance-based indicators, depending on the observation probability.

### Change in mosquito feeding preference for all the indicators and monitoring scenarios

**Figure S12.**
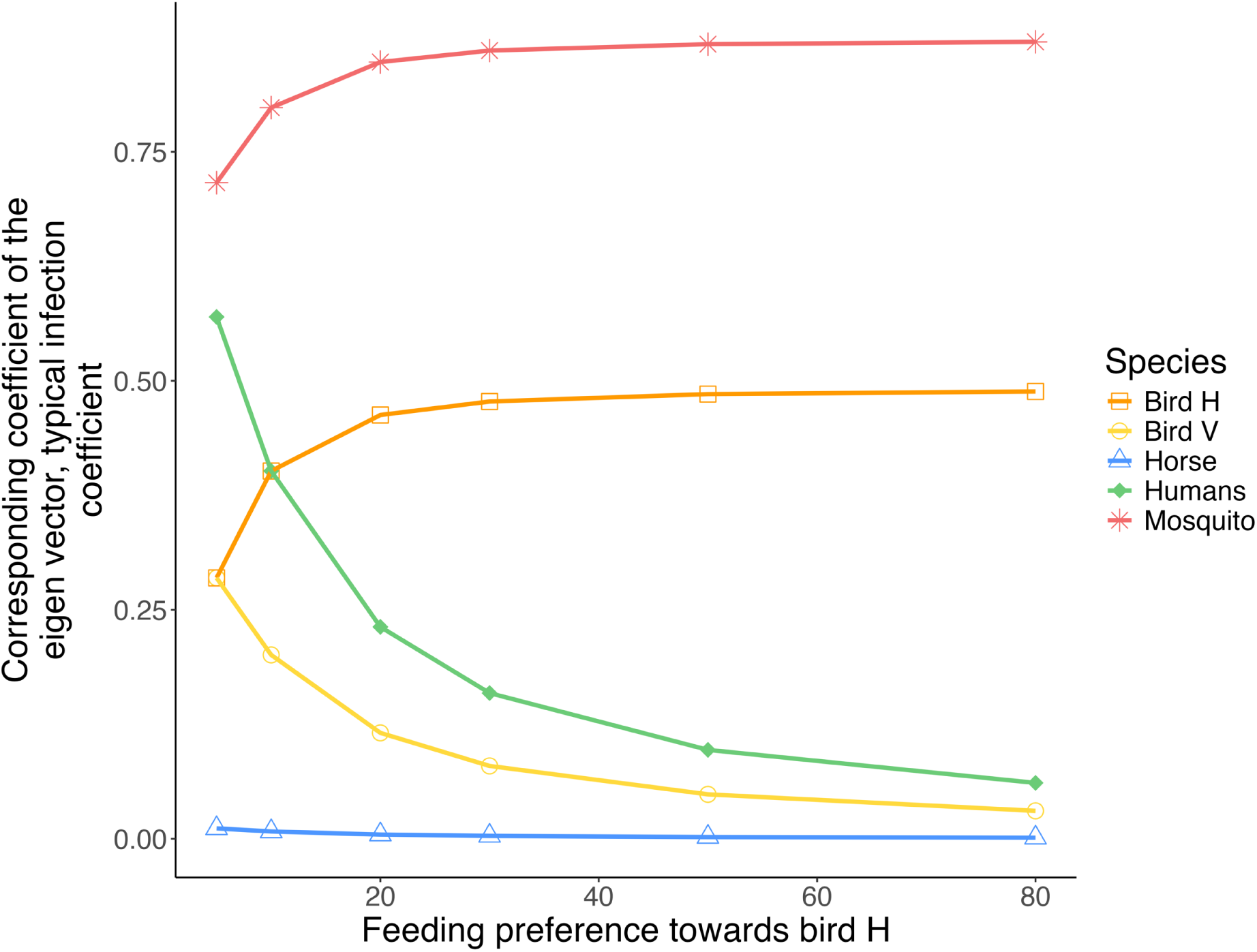
Relation between the feeding preference towards bird H and the *typical infection coefficient*, i.e. the corresponding coefficient of the eigenvector associated with the dominant eigenvalue for each species in the model.

**Figure S13.**
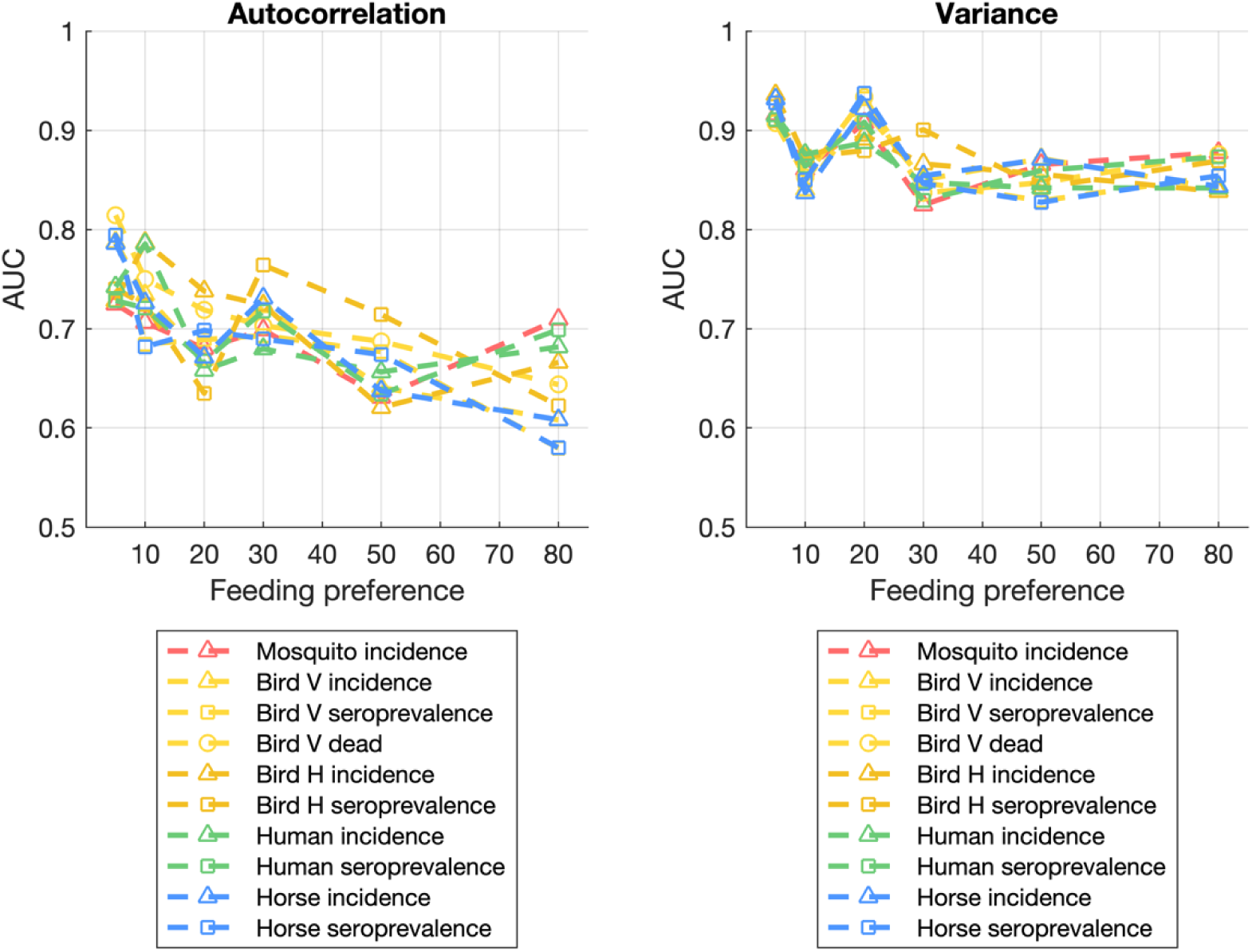
Prediction performance all univariate time series for both autocorrelation and variance, depending on the feeding preference towards bird H.

**Figure S14.**
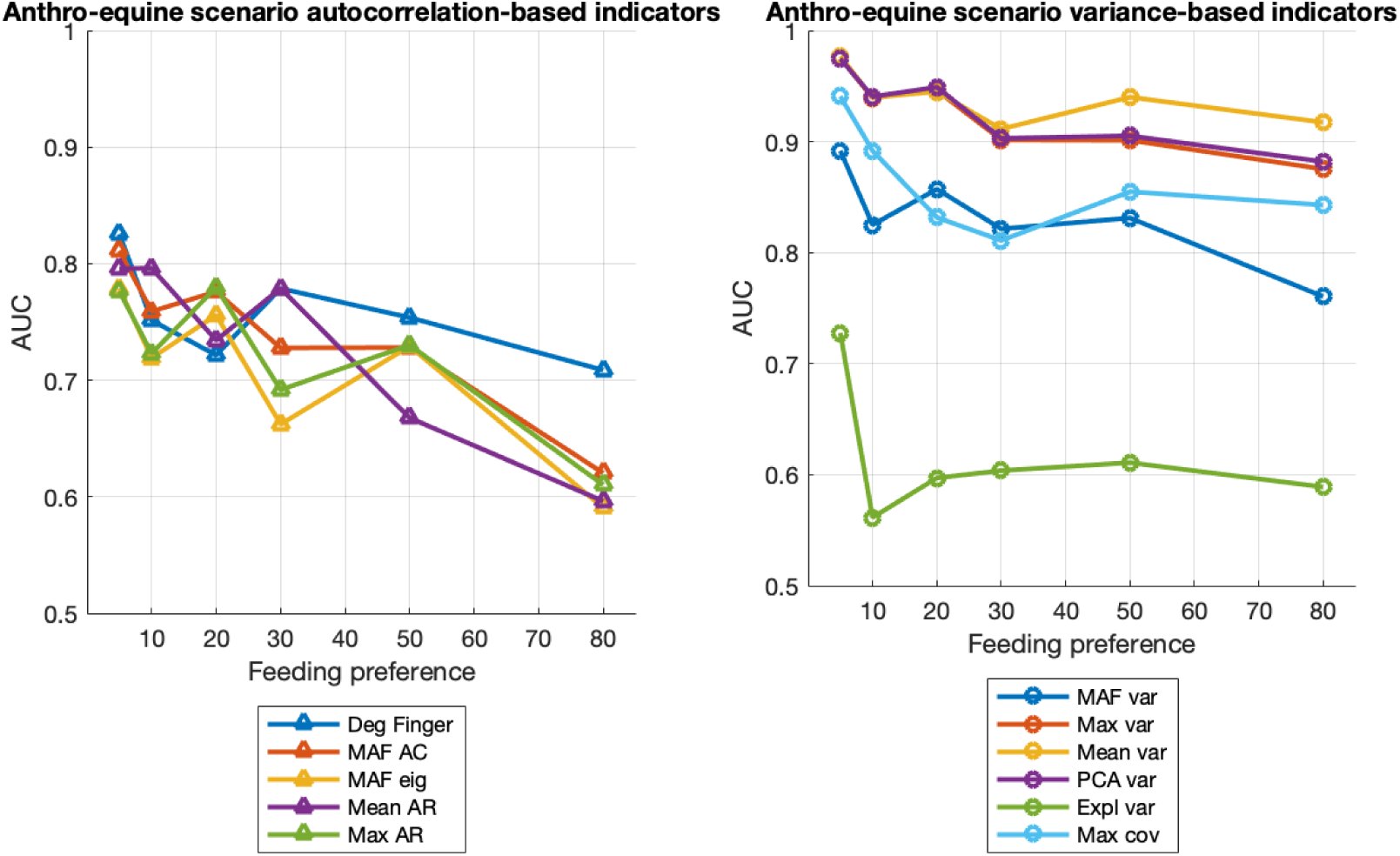
Prediction performance for the anthro-equine scenario for all autocorrelation and variance-based indicators, depending on the feeding preference towards bird H.

**Figure S15.**
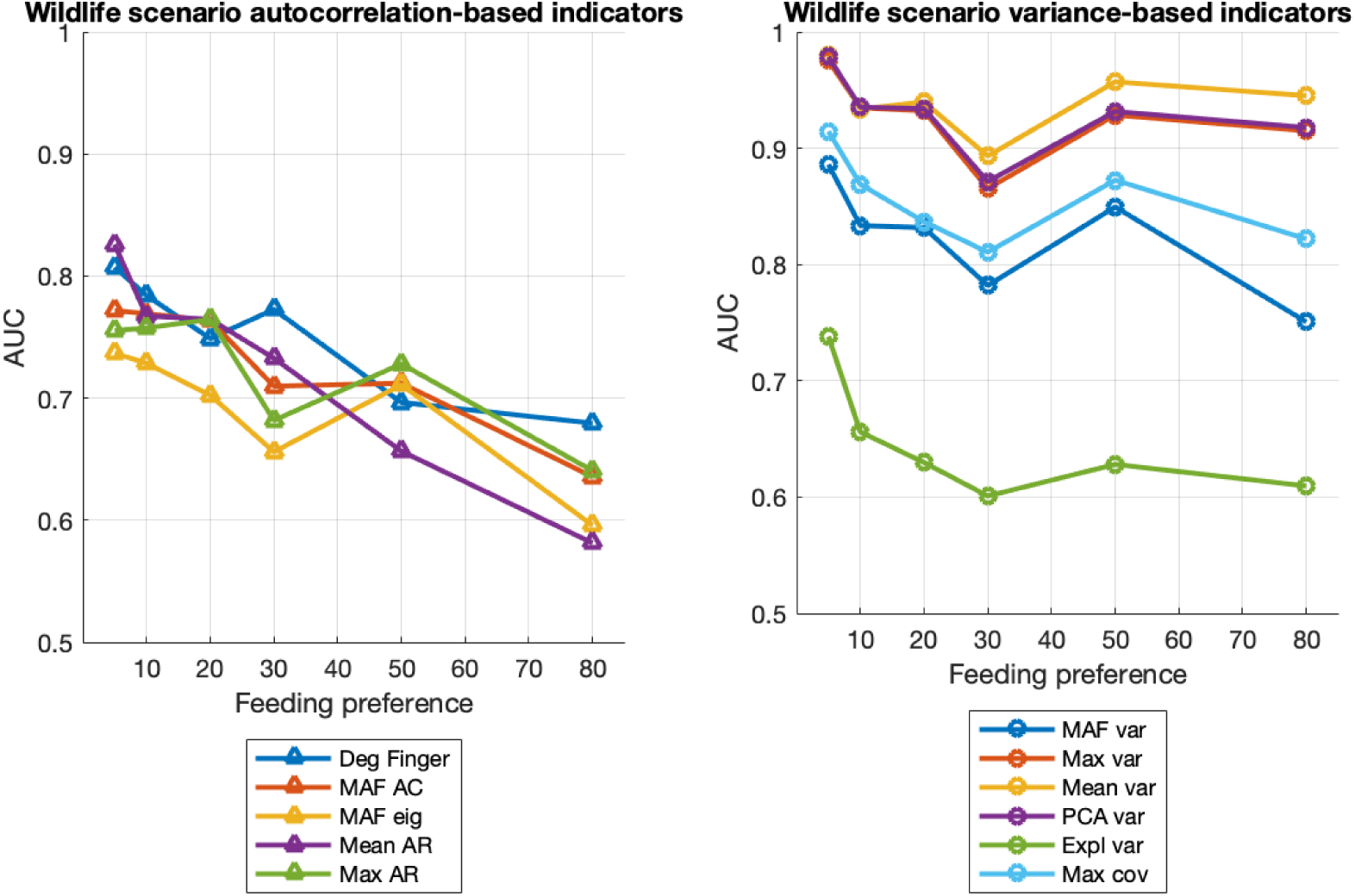
Prediction performance for the wildlife scenario for all autocorrelation and variance-based indicators, depending on the feeding preference towards bird H.

**Figure S16.**
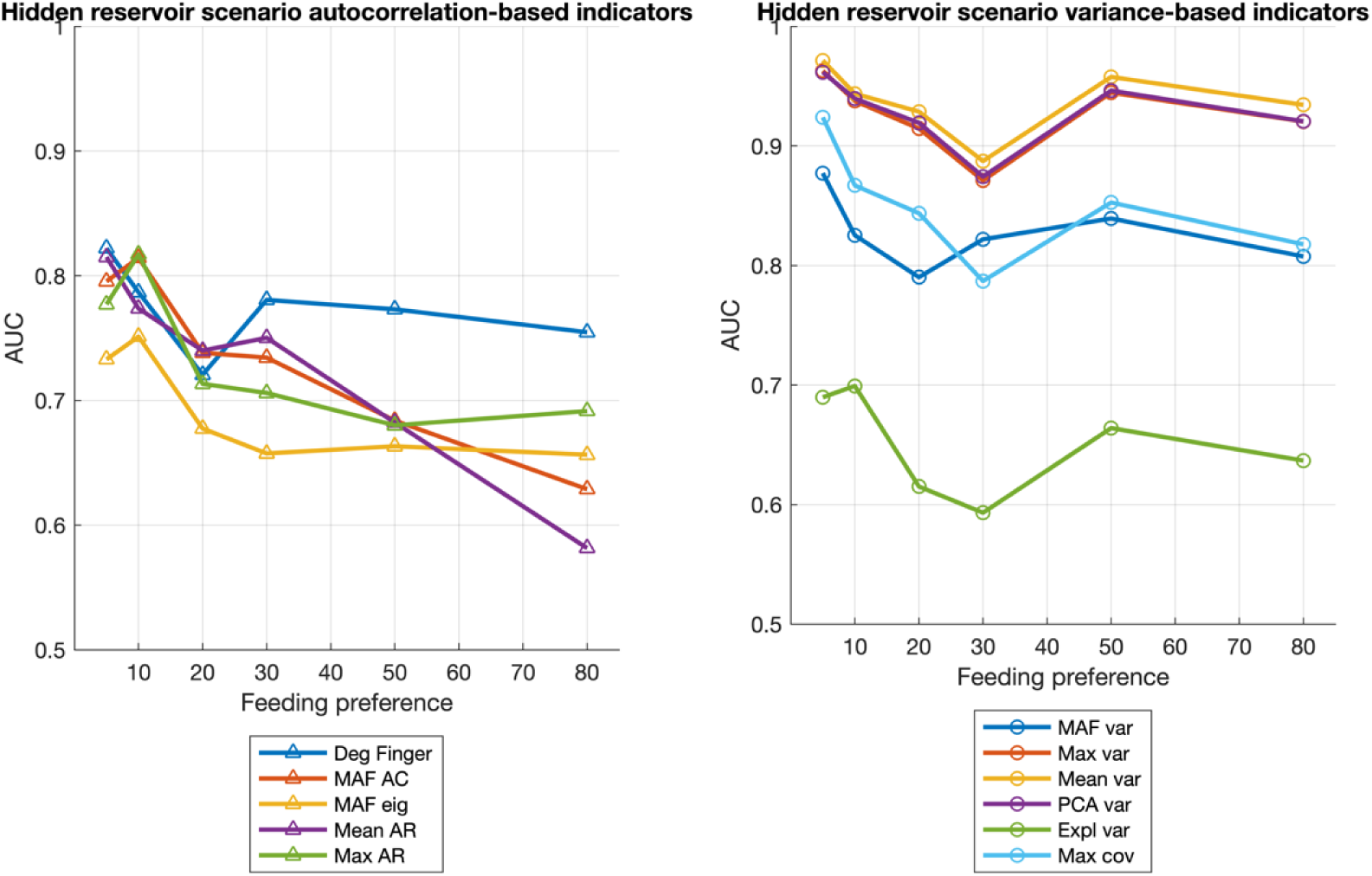
Prediction performance for the hidden reservoir scenario for all autocorrelation and variance-based indicators, depending on the feeding preference towards bird H.

**Figure S17.**
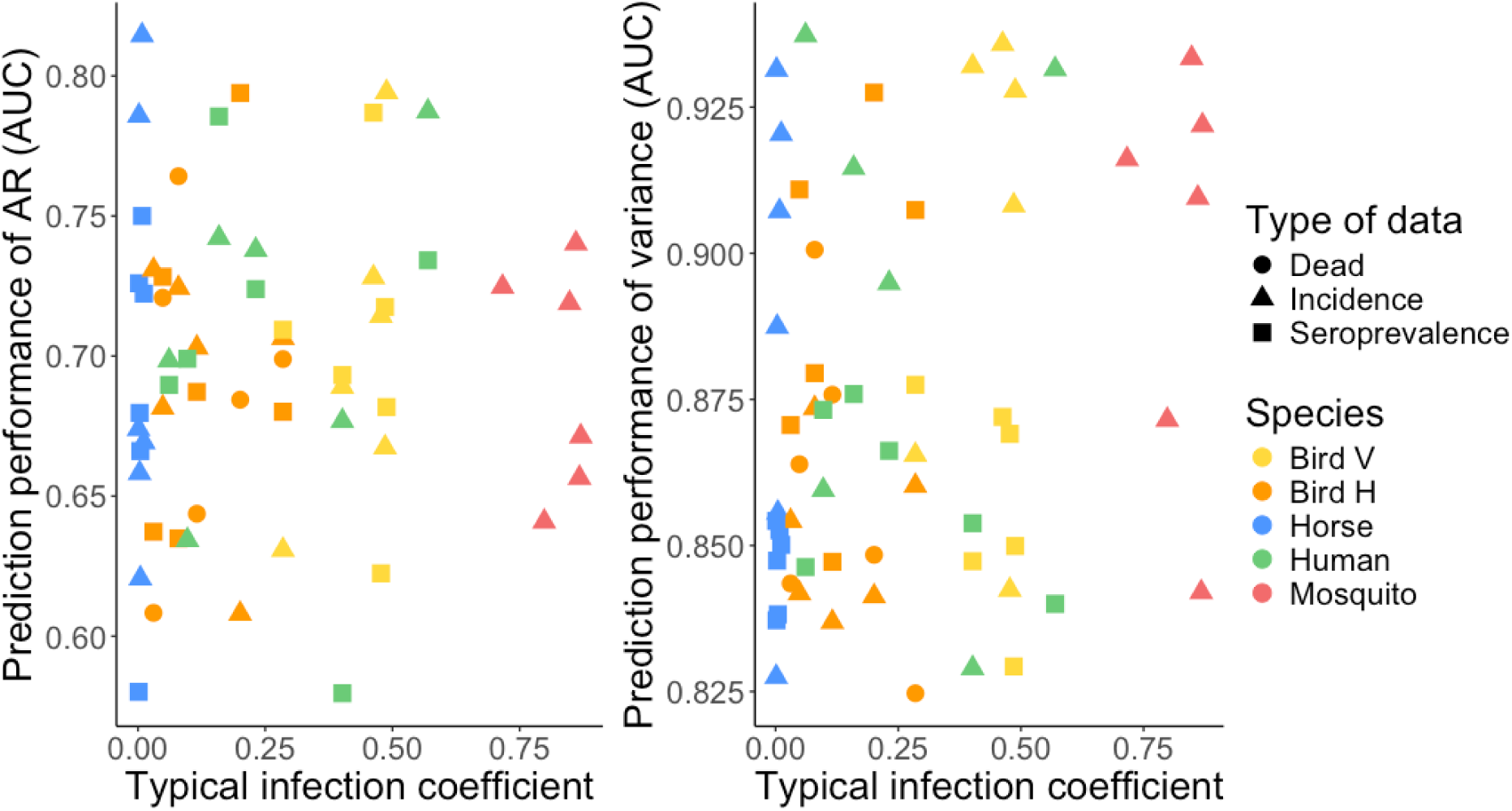
Prediction performance of univariate time series depending on the *typical infection coefficient*, i.e. the corresponding coefficient of the eigenvector associated with the dominant eigenvalue for each species in the model.

### Robustness of the results of different feeding preference for lower resolution and observation probability

The results of Figure S4 (reducing the resolution), and Figure S8 (reducing the observation probability) are reproduced for the most extreme feeding preference towards bird H (p_BH_=80) to test the robustness of the results of Figure S13 and verify that no variability in the prediction performance of the different species arises.

**Figure S18:**
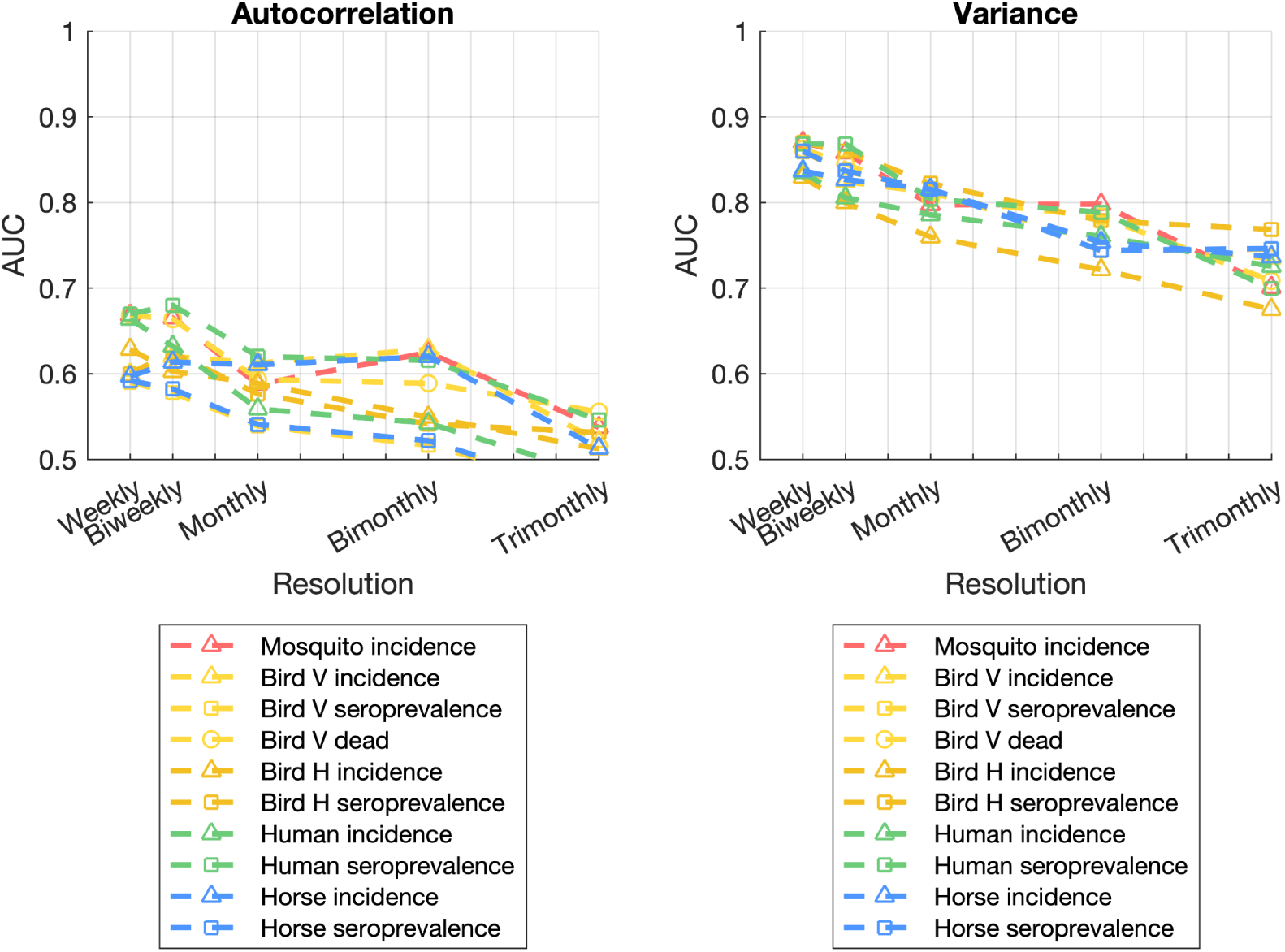
Prediction performance all univariate time series for both autocorrelation and variance, depending on the resolution of the data, for a strong feeding preference towards bird H (p_BH_=80).

**Figure S19:**
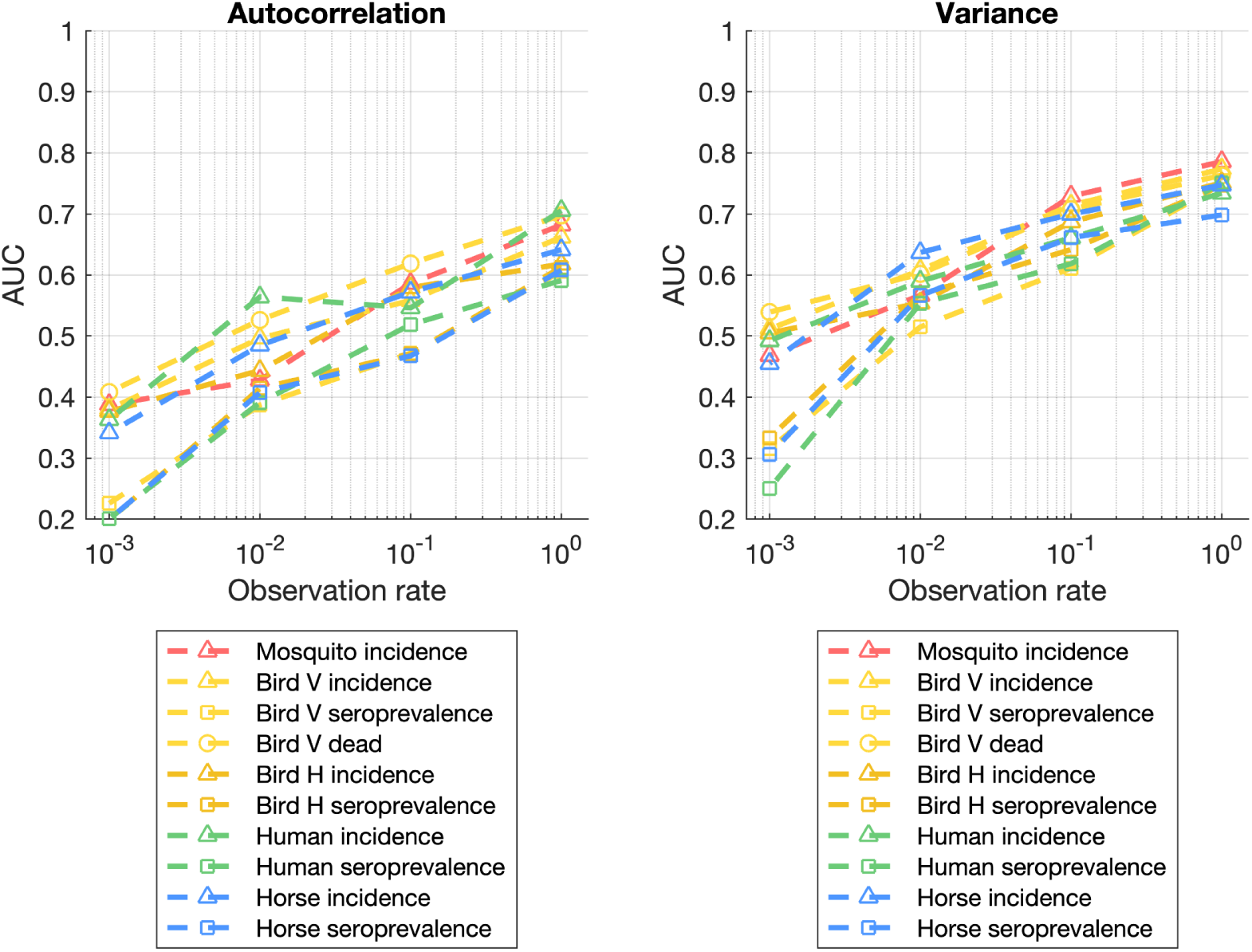
Prediction performance all univariate time series for both autocorrelation and variance, depending on the resolution of the data, for a strong feeding preference towards bird H (p_BH_=80).

### Change in bird relative abundances as a sensitivity analysis

To make sure that the effect of *the typical infection coefficient* on the prediction performance of a given species is not singular to the feeding preference coefficient, we reproduced the same analyses by varying the relative abundance of bird H compared to bird V and keeping the feeding preference coefficient at the default value. We generated time series of the model with 4 values for the relative abundance of bird species H. In the default emergence time series, bird H has the same abundance as bird species V (relative abundance *r_A_* = 1). Additionally, we generated time series with r_A_=0.75, 1.25, and 1.5. Similarly to the analyses performed by varying the feeding preference coefficient, for each simulation, we adapted the range of values for the biting rate to keep a range of R_0_ going from 0.7 to 1 for the emergence time series, and R_0_=0.8 for the stable time series.

**Figure S20.**
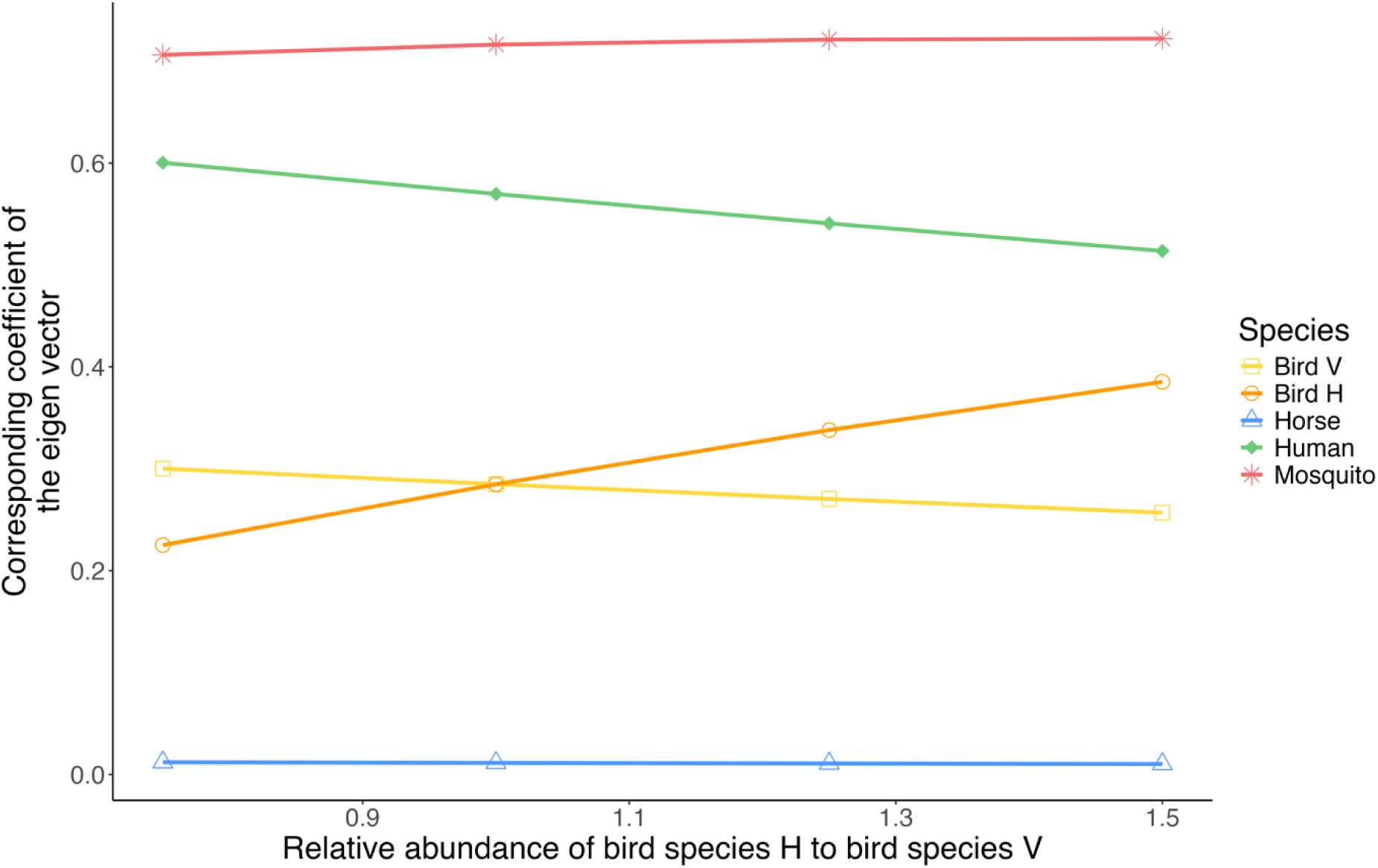
Relation between the relative abundance of bird H and the *typical infection coefficient*, i.e. the corresponding coefficient of the eigenvector associated with the dominant eigenvalue for each species in the model.

**Figure S21.**
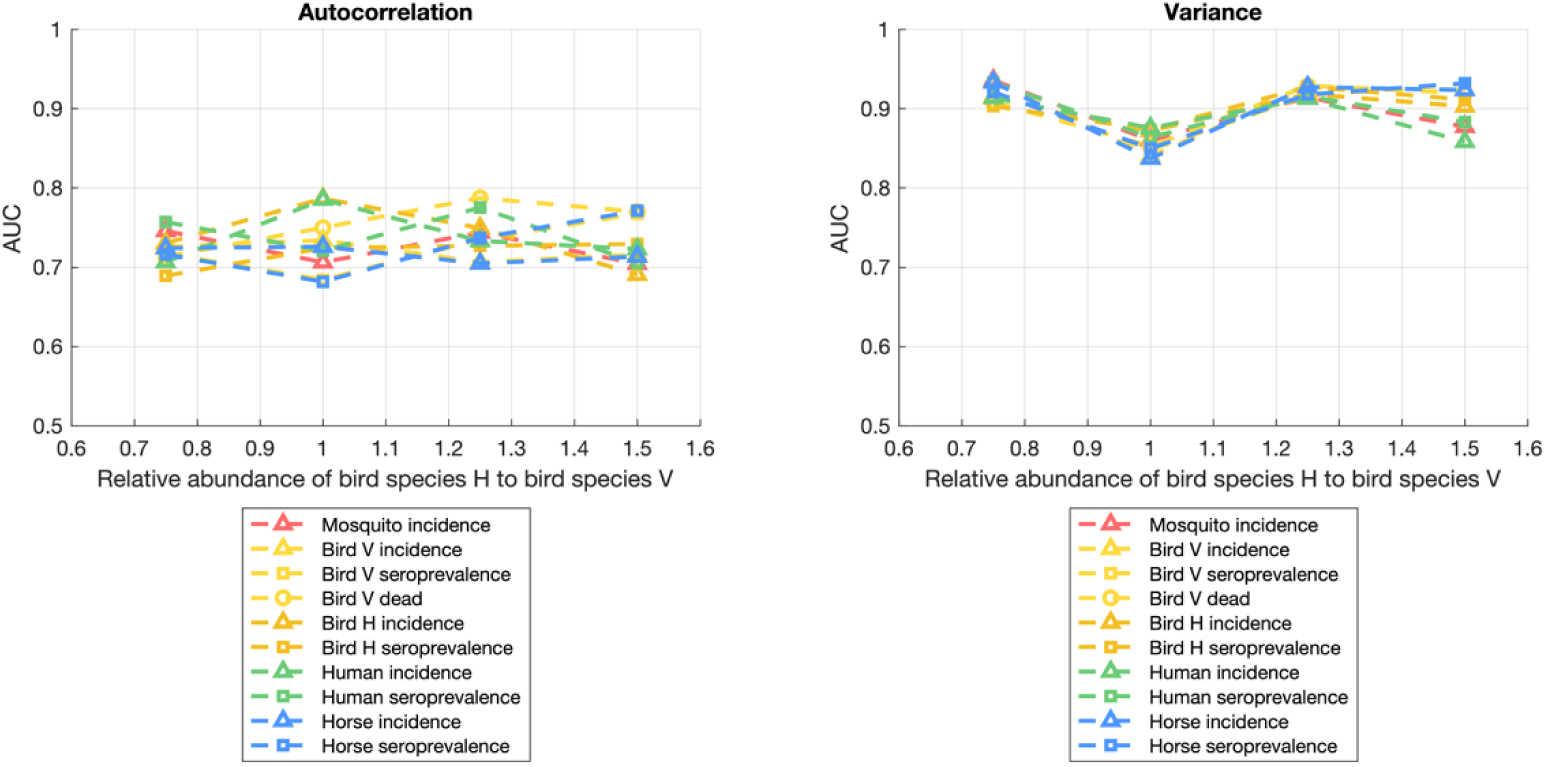
Prediction performance all univariate time series for both autocorrelation and variance, depending on the relative abundance of bird H.

**Figure S22.**
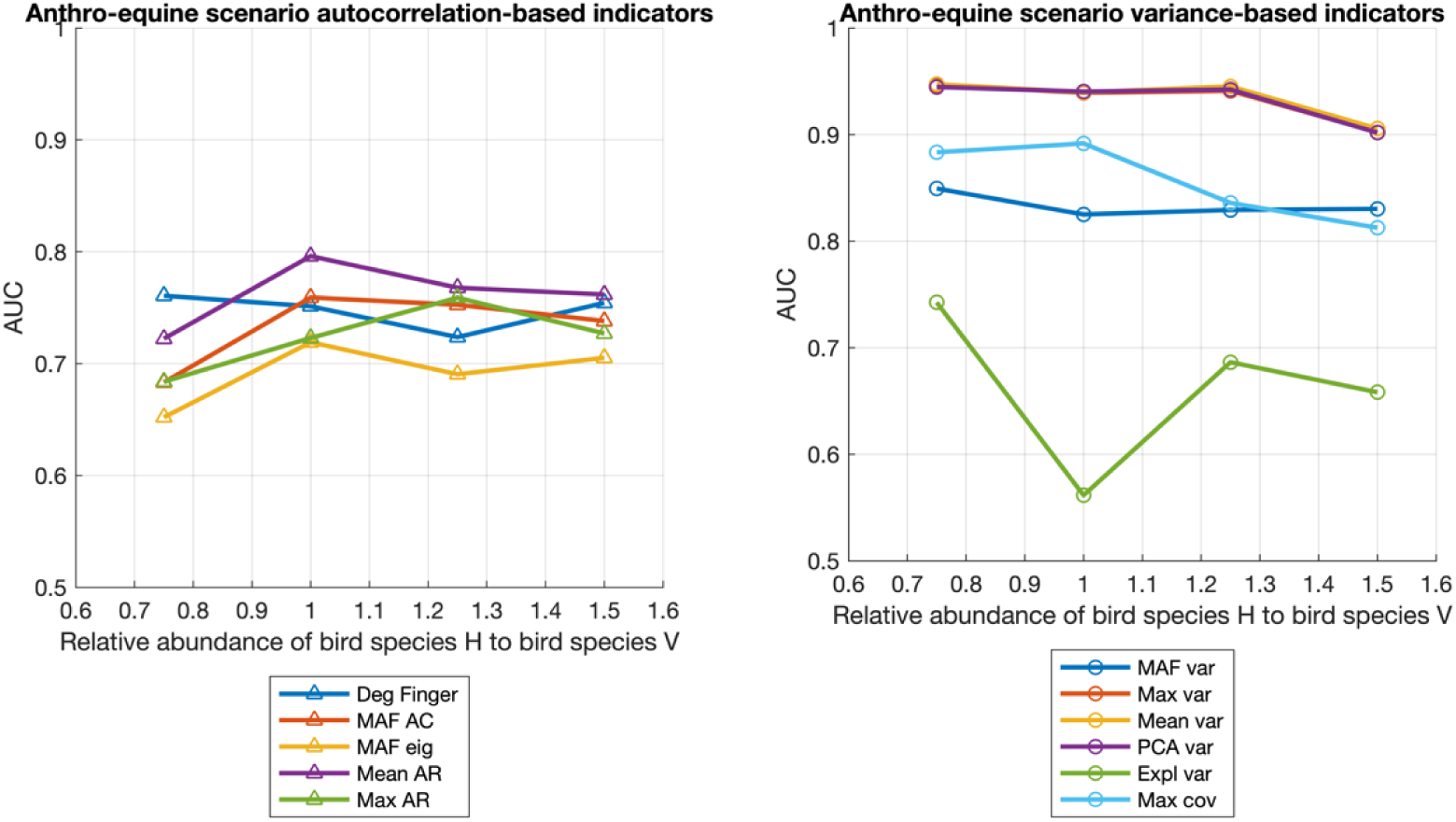
Prediction performance for the anthro-equine scenario for all autocorrelation and variance-based indicators, depending on the relative abundance of bird H.

**Figure S23.**
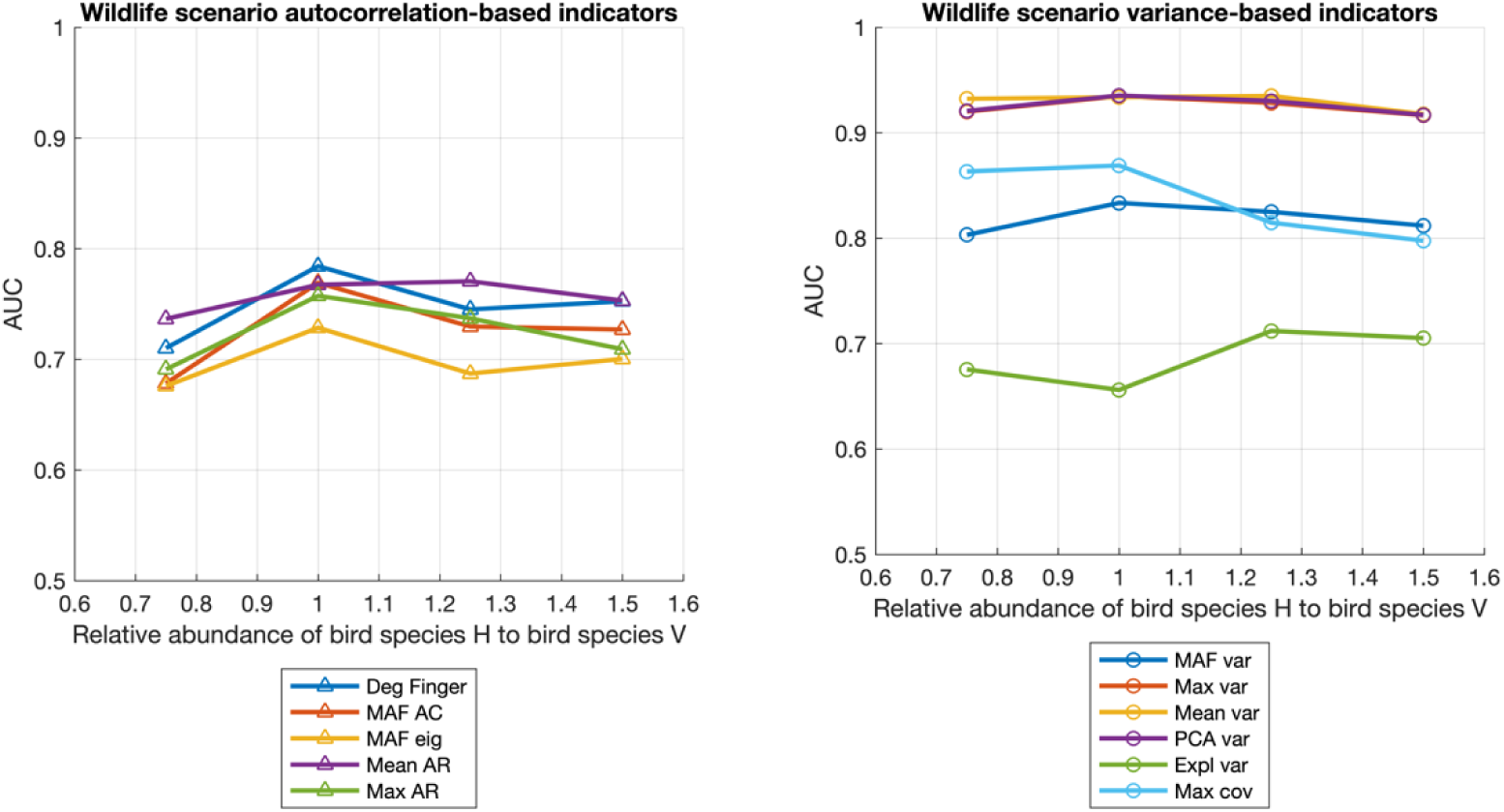
Prediction performance for the wildlife scenario for all autocorrelation and variance-based indicators, depending on the relative abundance of bird H.

**Figure S24.**
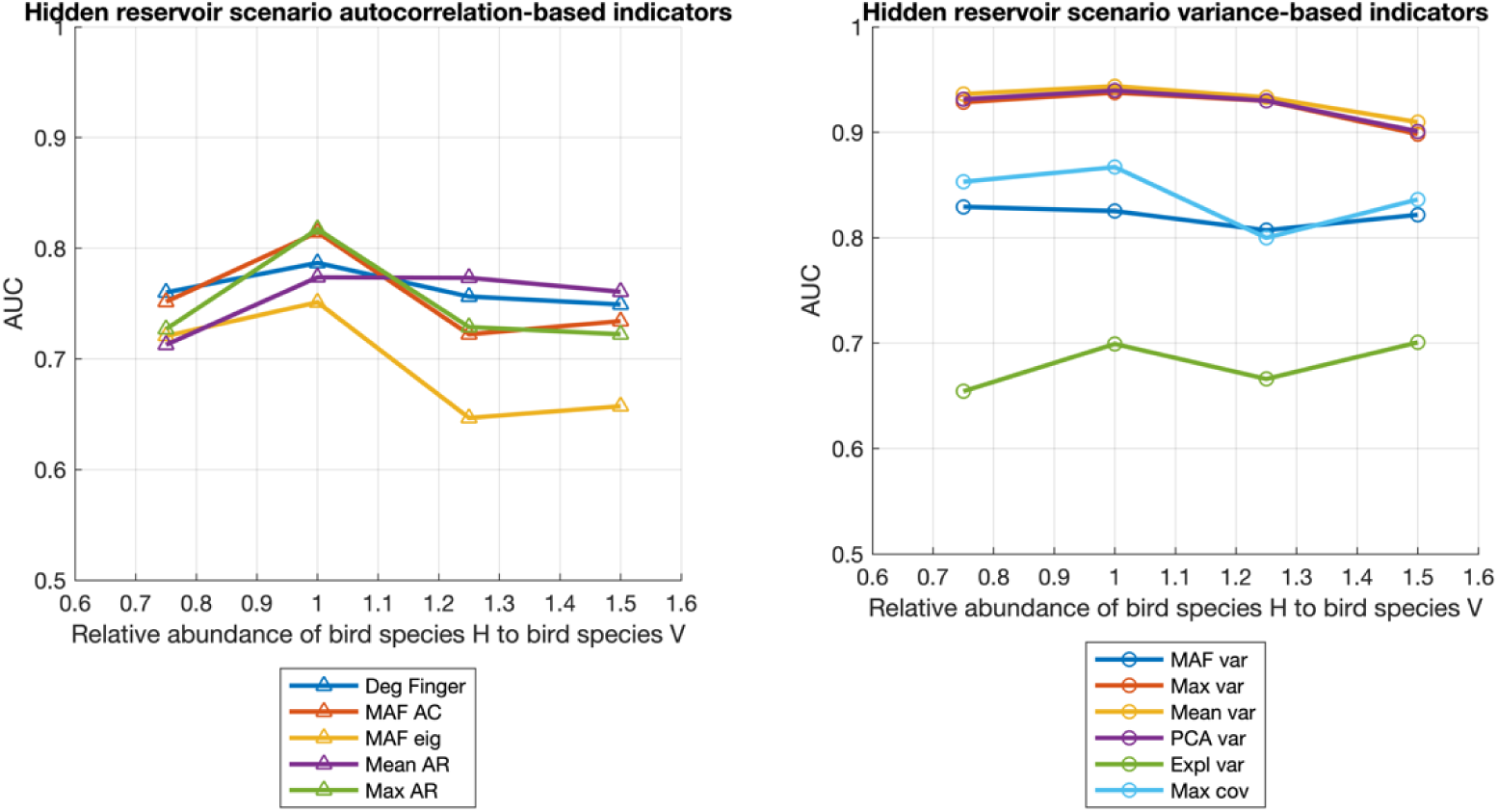
Prediction performance for the hidden reservoir scenario for all autocorrelation and variance-based indicators, depending on the relative abundance of bird H.

**Figure S25.**
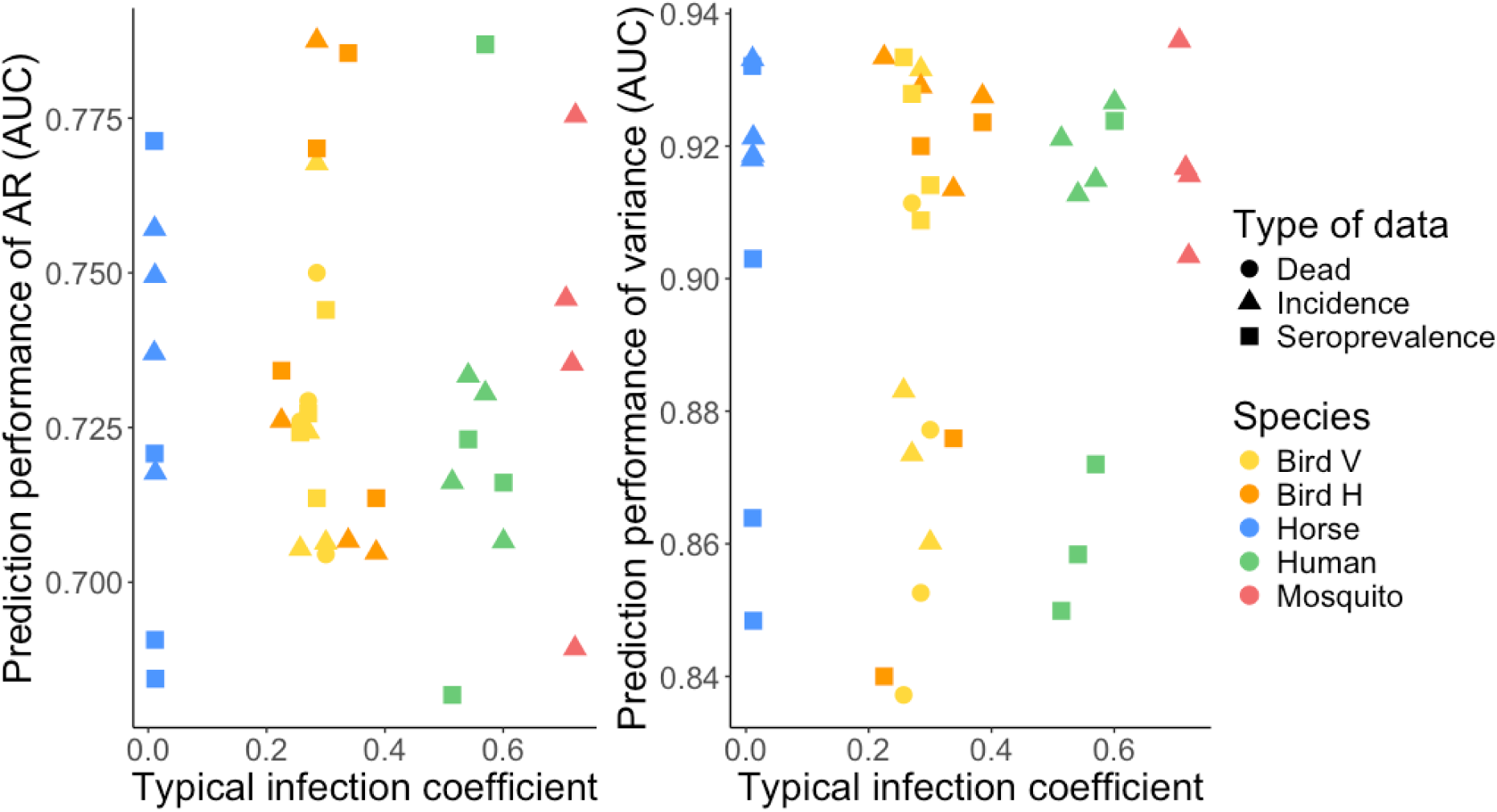
Prediction performance of univariate time series depending on the *typical infection coefficient*, i.e. the corresponding coefficient of the eigenvector associated with the dominant eigenvalue for each species in the model, calculated by varying the relative abundance of bird species H.

### Lead time of prediction

**Figure S26.**
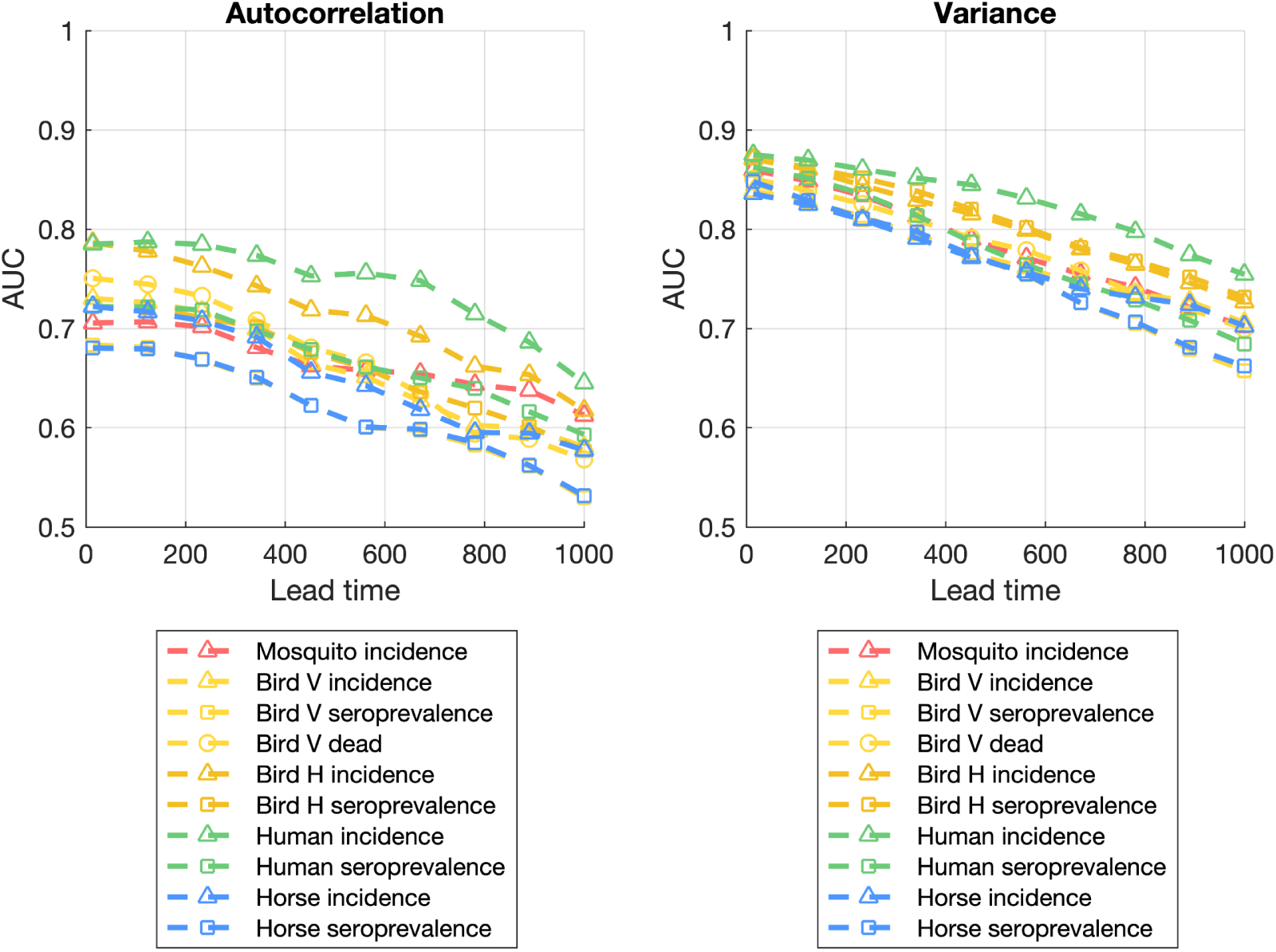
Prediction performance all univariate time series for both autocorrelation and variance, for different lead times (days).

**Figure S27.**
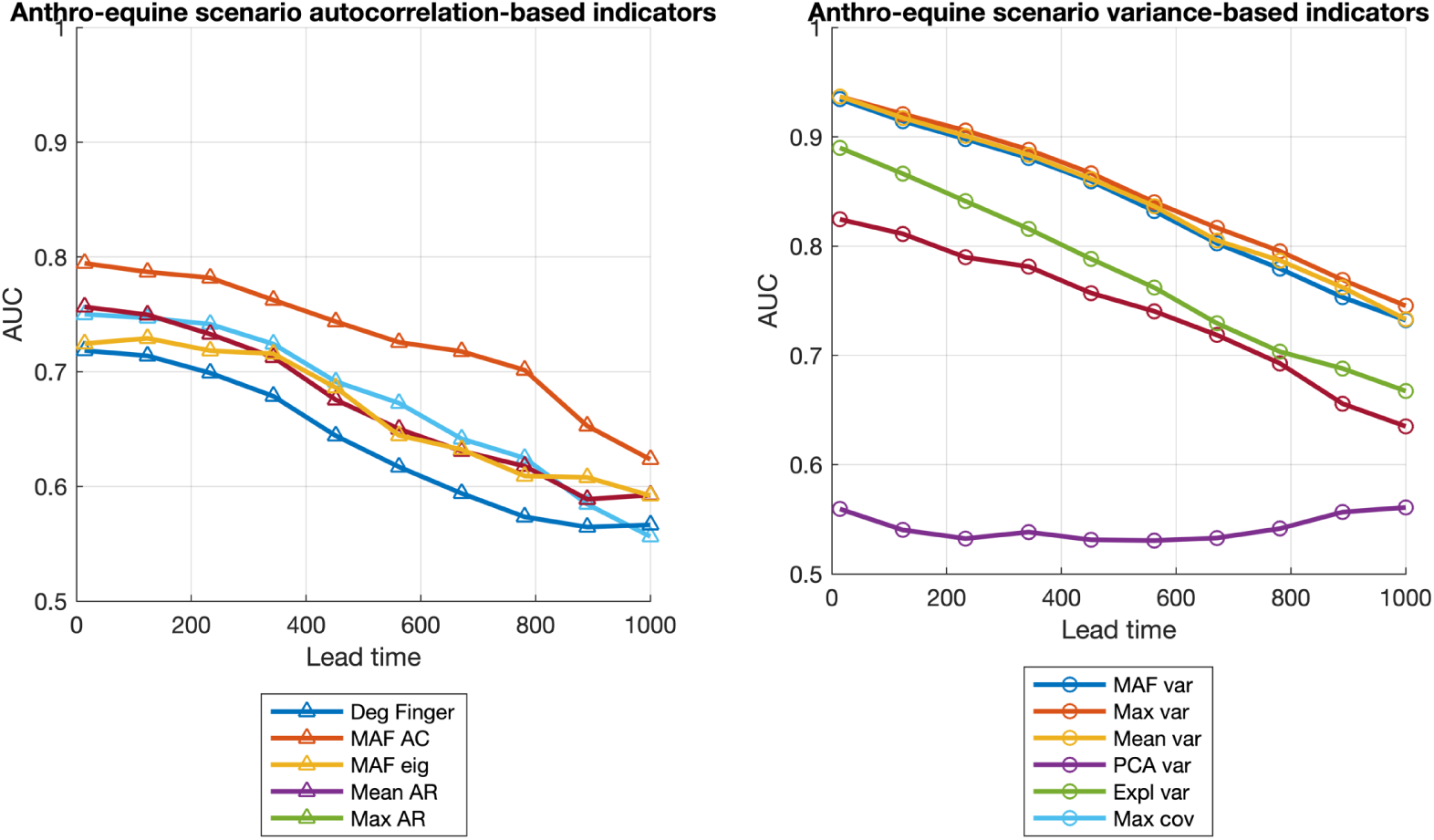
Prediction performance for the Anthro-equine scenario for all autocorrelation and variance-based indicators, for different lead times (days).

**Figure S28.**
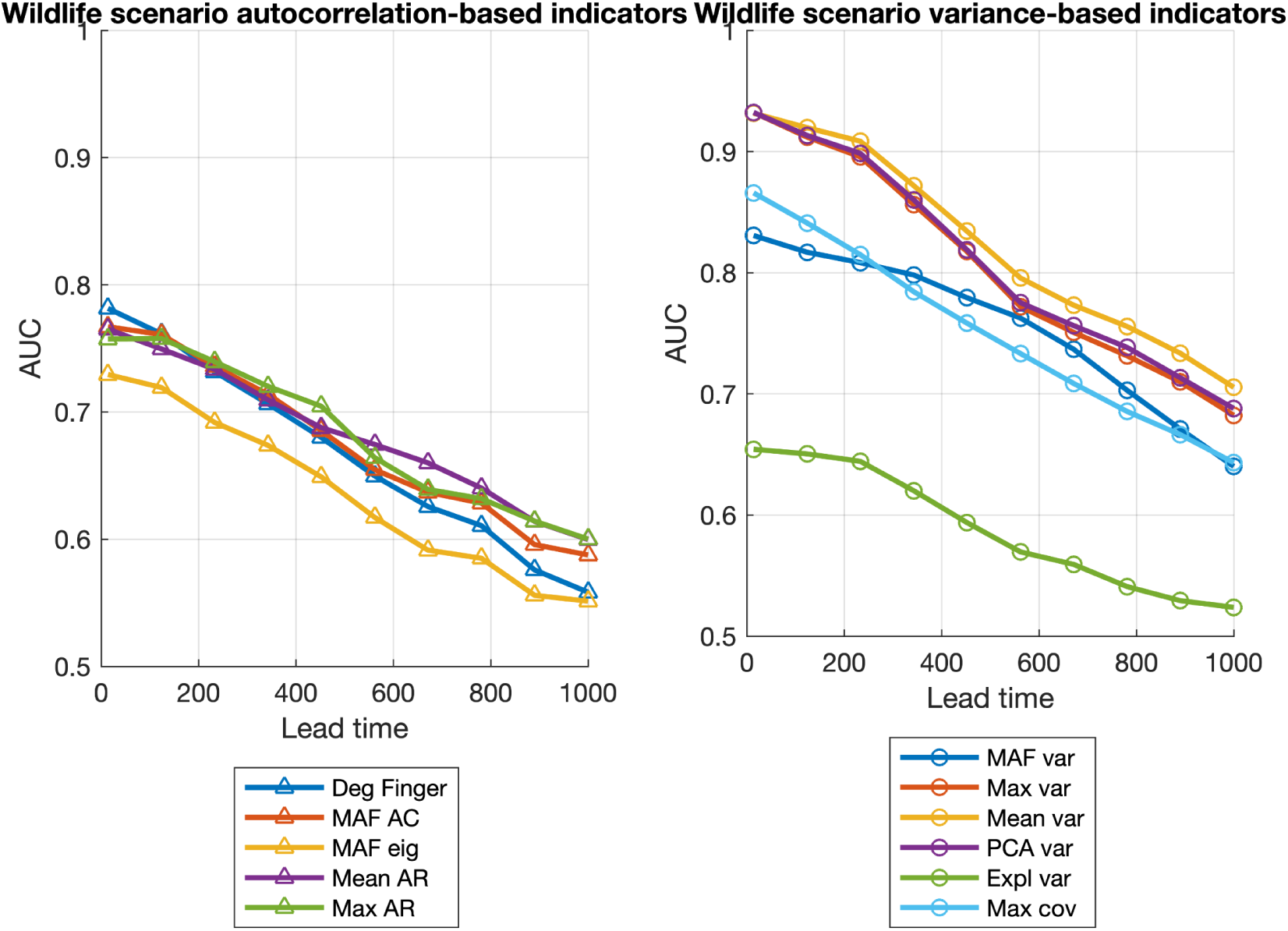
Prediction performance for the Wildlife scenario for all autocorrelation and variance-based indicators, for different lead times (days).

**Figure S29.**
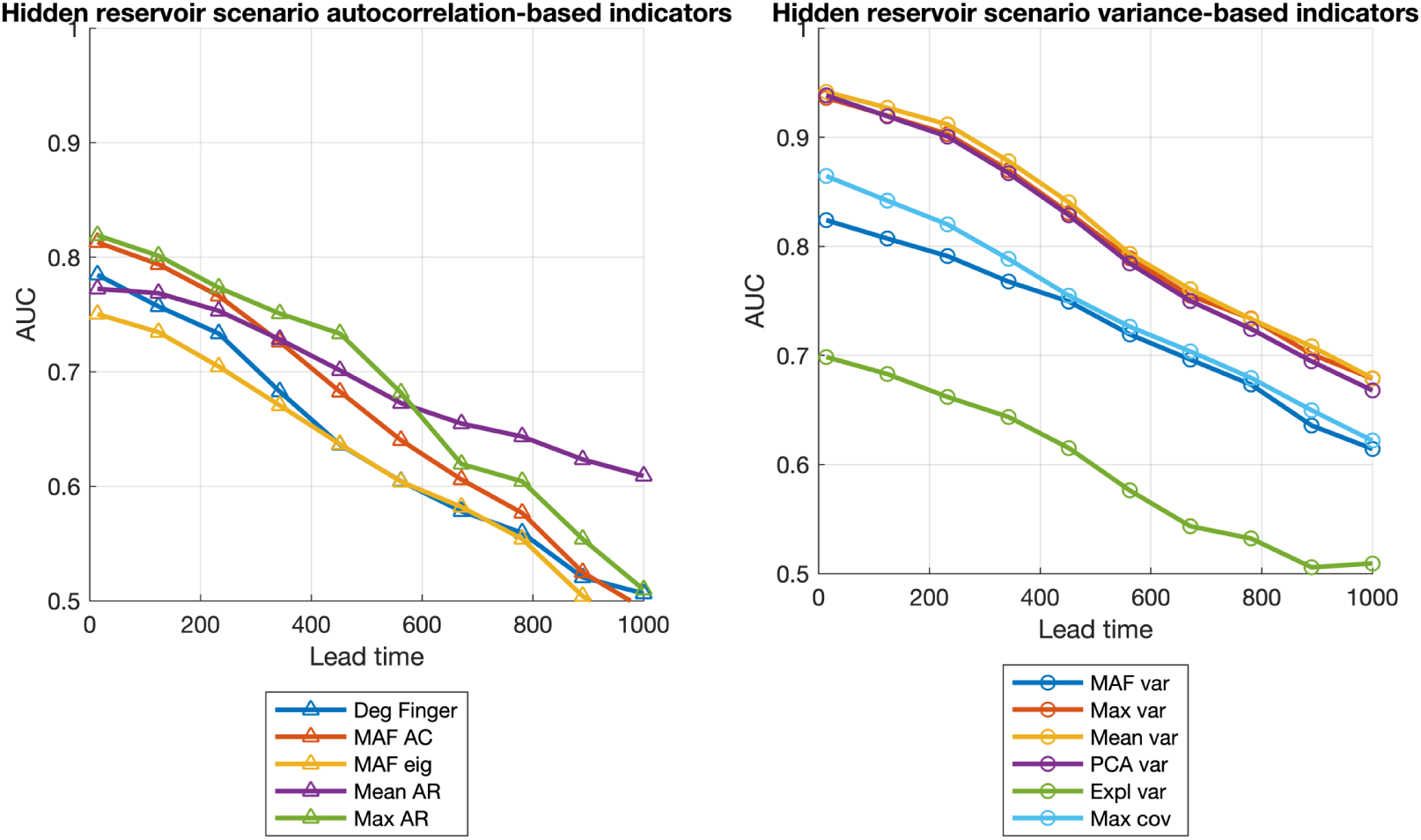
Prediction performance for the hidden reservoir scenario for all autocorrelation and variance-based indicators, for different lead times (days).

## Notes

### Competing Interest Statement

The authors have declared no competing interest.

